# *Stxbp1/Munc18-1* haploinsufficiency in mice recapitulates key features of *STXBP1* encephalopathy and impairs cortical inhibition

**DOI:** 10.1101/621516

**Authors:** Wu Chen, Zhao-Lin Cai, Eugene S. Chao, Hongmei Chen, Shuang Hao, Hsiao-Tuan Chao, Joo Hyun Kim, Jessica E. Messier, Huda Y. Zoghbi, Jianrong Tang, John W. Swann, Mingshan Xue

## Abstract

Mutations in genes encoding synaptic proteins cause many neurodevelopmental disorders, but the underlying pathogeneses are poorly understood. Syntaxin-binding protein 1 (STXBP1) is an essential component of the neurotransmitter release machinery. Its *de novo* heterozygous mutations are among the most frequent causes of neurodevelopmental disorders including intellectual disabilities and epilepsies. These disorders, collectively referred to as *STXBP1* encephalopathy, affect a broad spectrum of neurological and neuropsychiatric features common among neurodevelopmental disorders. To gain insight into *STXBP1* encephalopathy pathogenesis, we generated new *Stxbp1* null alleles in mice and found that *Stxbp1* haploinsufficiency impaired cognitive, psychiatric, and motor functions and caused cortical hyperexcitability and seizures. Surprisingly, *Stxbp1* haploinsufficiency reduced neurotransmission from cortical parvalbumin- and somatostatin-expressing GABAergic interneurons by differentially decreasing the synaptic strength and connectivity, respectively. These results demonstrate that *Stxbp1* haploinsufficient mice recapitulate key features of *STXBP1* encephalopathy and indicate that inhibitory dysfunction is likely a key contributor to the disease pathogenesis.

## Introduction

Human genetic studies of neurodevelopmental disorders continue to uncover pathogenic variants in genes encoding synaptic proteins (Deciphering Developmental Disorders Study, 2017; 2015; Hoischen et al., 2014; Lindy et al., 2018; Stessman et al., 2017; Zhu et al., 2014), demonstrating the importance of these proteins for neurological and neuropsychiatric features. The molecular and cellular functions of many of these synaptic proteins have been extensively studied.

However, to understand the pathological mechanisms underlying these synaptic disorders, in-depth neurological and behavioral studies in animal models are necessary. This knowledge gap can be significantly narrowed by studying a few prioritized genes that are highly penetrant and affect a broad spectrum of neurological and neuropsychiatric features common among neurodevelopmental disorders (Hoischen et al., 2014; Ogden et al., 2016). Syntaxin-binding protein 1 (STXBP1, also known as MUNC18-1) is one such example because its molecular and cellular functions are well understood (Rizo and Xu, 2015), its mutations are emerging as prevalent causes of multiple neurodevelopmental disorders (Stamberger et al., 2016), and yet it remains unclear how its dysfunction causes diseases.

Stxbp1/Munc18-1 is involved in synaptic vesicle docking, priming, and fusion through multiple interactions with the neuronal soluble *N*-ethylmaleimide-sensitive factor-attachment protein receptors (SNAREs) (Rizo and Xu, 2015). Genetic deletion of Stxbp1 in worms, flies, mice, and fish abolishes neurotransmitter release and leads to lethality and cell-intrinsic degeneration of neurons (Grone et al., 2016; Harrison et al., 1994; Heeroma et al., 2004; Verhage et al., 2000; Weimer et al., 2003). In humans, *STXBP1 de novo* heterozygous mutations cause several of the most severe forms of epileptic encephalopathies including Ohtahara syndrome (Saitsu et al., 2008; 2010), West syndrome (Deprez et al., 2010; Otsuka et al., 2010), Lennox-Gastaut syndrome (Carvill et al., 2013; Epi4K Consortium et al., 2013), Dravet syndrome (Carvill et al., 2014), and other types of early-onset epileptic encephalopathies (Deprez et al., 2010; Mignot et al., 2011; Stamberger et al., 2016). Furthermore, *STXBP1* is one of the most frequently mutated genes in sporadic intellectual disabilities and developmental disorders (Deciphering Developmental Disorders Study, 2017; 2015; Hamdan et al., 2011; 2009; Rauch et al., 2012; Suri et al., 2017). All *STXBP1* encephalopathy patients show intellectual disability, mostly severe to profound, and 95% of patients have epilepsy (Stamberger et al., 2016). Other clinical features that are present in subsets of patients include developmental delay, dystonia, ataxia, hypotonia, tremor, hyperactivity, anxiety, stereotypies, aggressive behaviors, and autistic features (Boutry-Kryza et al., 2015; Campbell et al., 2012; Deprez et al., 2010; Hamdan et al., 2009; Mignot et al., 2011; Milh et al., 2011; Rauch et al., 2012; Stamberger et al., 2016; Suri et al., 2017; Weckhuysen et al., 2013).

*STXBP1* encephalopathy is mostly caused by haploinsufficiency because more than 60% of the reported mutations are deletions and nonsense, frameshift, or splice site variants (Stamberger et al., 2016). A subset of missense variants were shown to destabilize the protein (Guiberson et al., 2018; Kovačević et al., 2018; Saitsu et al., 2010; 2008) and act as dominant negatives to further reduce the wild type protein levels (Guiberson et al., 2018). Thus, partial loss-of-function of *Stxbp1 in vivo* would offer opportunities to model *STXBP1* encephalopathy and study its pathogenesis. Indeed, removing *stxbp1b*, one of the two *STXBP1* homologs in zebrafish, caused spontaneous electrographic seizures (Grone et al., 2016). Three different *Stxbp1* null alleles have been generated in mice (Kovačević et al., 2018; Miyamoto et al., 2017; Verhage et al., 2000). However, characterizations of the corresponding heterozygous knockout mice were limited and yielded inconsistent results. For example, the reported cognitive phenotypes in mutant mice are mild or inconsistent between studies (Kovačević et al., 2018; Miyamoto et al., 2017; Orock et al., 2018), whereas human patients usually have severe intellectual disability. Seizures were observed in one study (Kovačević et al., 2018), but not in another using the same line of mutant mice (Orock et al., 2018). Motor and a number of neuropsychiatric dysfunctions were not reported in previous studies (Hager et al., 2014; Kovačević et al., 2018; Miyamoto et al., 2017; Orock et al., 2018). Thus, a comprehensive neurological and behavioral study of *Stxbp1* haploinsufficiency models is still lacking and it is also unclear to what extent *Stxbp1* haploinsufficient mice can recapitulate the neurological and neuropsychiatric phenotypes of *STXBP1* encephalopathy. More importantly, it remains elusive how *STXBP1* haploinsufficiency *in vivo* leads to hyperexcitable neural circuits and neurological deficits. To address these questions, we developed two new *Stxbp1* haploinsufficiency mouse models and found that they recapitulated all key phenotypes of human patients, including impaired cognitive, psychiatric, and motor functions and seizures. Electrophysiological experiments in *Stxbp1* haploinsufficient mice revealed a reduction of GABAergic synaptic transmission via different mechanisms from two main classes of cortical inhibitory neurons, parvalbumin-expressing (Pv) and somatostatin-expressing (Sst) interneurons. Thus, these results demonstrate a crucial role of *Stxbp1* in neurological and neuropsychiatric functions and indicate that *Stxbp1* haploinsufficient mice are construct and face valid models of *STXBP1* encephalopathy. The reduced inhibition is likely a major contributor to the cortical hyperexcitability and neurobehavioral phenotypes of *Stxbp1* haploinsufficient mice. The differential effects on Pv and Sst interneuron-mediated inhibition also suggest synapse-specific functions of *Stxbp1* in neural circuits.

## Results

### Generation of two new *Stxbp1* null alleles

To model *STXBP1* haploinsufficiency in mice, we first generated a knockout-first (KO-first) allele (*tm1a*), in which *Stxbp1* genomic locus was targeted with a multipurpose cassette (Skarnes et al., 2011; Testa et al., 2004). The targeted allele contains a splice acceptor site from *Engrailed 2* (*En2SA*), an encephalomyocarditis virus internal ribosomal entry site (*IRES*), *lacZ*, and SV40 polyadenylation element (pA) that trap the transcripts after exon 6, thereby truncating the *Stxbp1* mRNA. The trapping cassette (*En2SA-IRES-lacZ-pA*) and exon 7 are flanked by two *FRT* sites and two *loxP* sites, respectively (***Figure 1-supplement 1A***). By sequentially crossing with Flp and Cre germline deleter mice, we removed both trapping cassette and exon 7 from heterozygous KO-first mice, which leads to a premature stop codon in exon 8, generating a conventional knockout (KO) allele (*tm1d*) (***Figure 1A***). Heterozygous KO (*Stxbp1^tm1d/+^*) and KO-first (*Stxbp1^tm1a/+^*) mice are maintained on the C57BL/6J isogenic background for all experiments.

**Figure 1.**
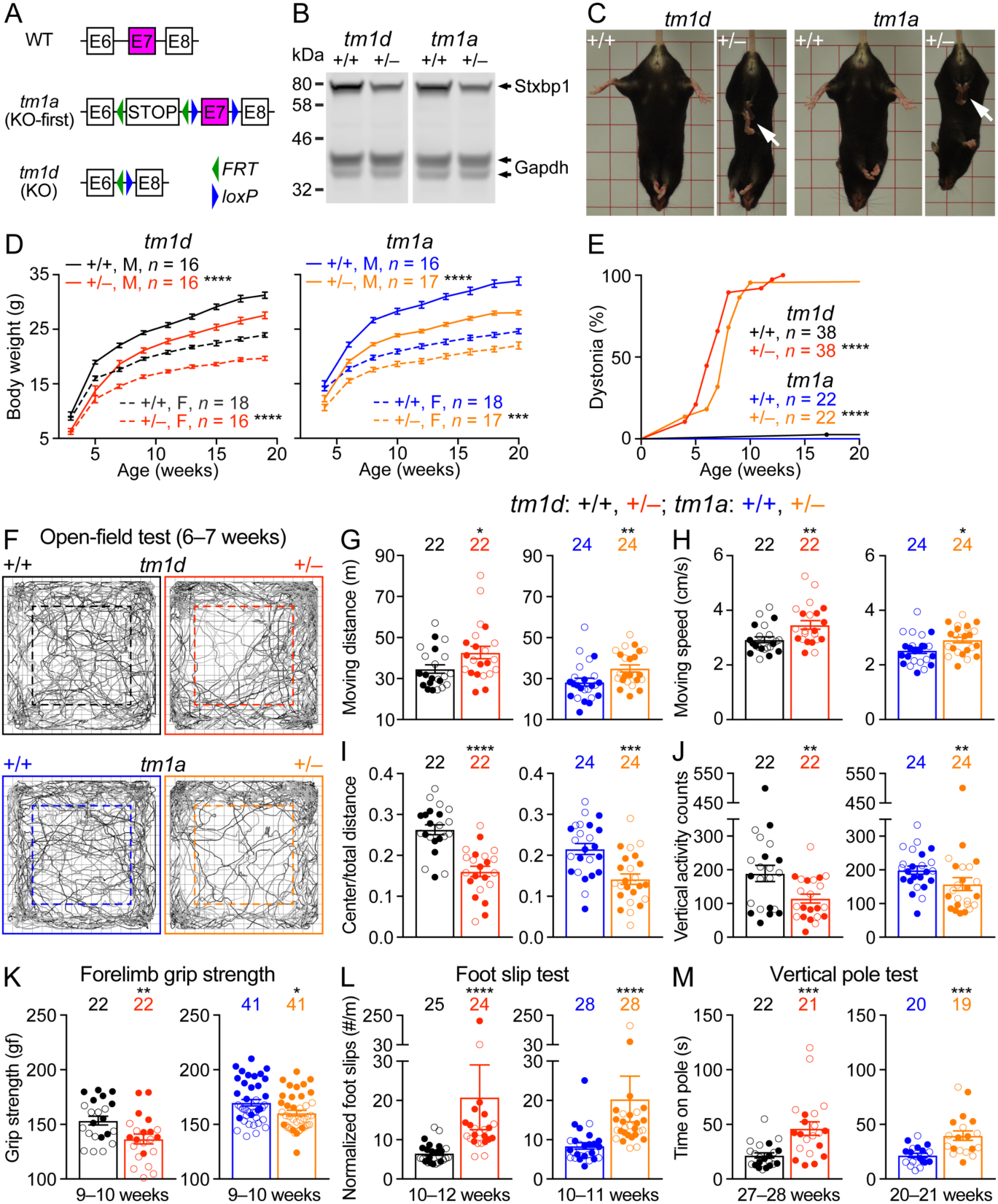
*Stxbp1* haploinsufficient mice exhibit dystonia, decreased body weights, and motor dysfunction. (**A**) Genomic structures of *Stxbp1* WT, *tm1a* (KO-first), and *tm1d* (KO) alleles. In the *tm1a* allele, the STOP including the *En2SA-lacZ-pA* trapping cassette (see ***Figure 1-supplement 1A***) truncates the *Stxbp1* mRNA after exon 6. In the *tm1d* allele, exon 7 is deleted, resulting in a premature stop codon in exon 8. E, exon; *FRT*, Flp recombination site; *loxP*, Cre recombination site. (**B**) Representative Western blots of proteins from the cortices of 3-month-old WT, *Stxbp1^tm1d/+^*, and *Stxbp1^tm1a/+^* mice. Gapdh, a housekeeping protein as loading control. Note reduced Stxbp1 levels in *Stxbp1^tm1d/+^* and *Stxbp1^tm1a/+^* mice. (**C**) *Stxbp1^tm1d/+^* and *Stxbp1^tm1a/+^* mice were smaller and showed dystonia and hindlimb clasping (arrows). (**D**) Body weights as a function of age. M, male; F, female. (**E**) The fraction of mice with dystonia as a function of age. (**F**) Representative tracking plots of the mouse positions in the open-field test. Note that *Stxbp1^tm1d/+^* and *Stxbp1^tm1a/+^* mice traveled less in the center (dashed box) than WT mice. (**G**–**J**) Summary data showing hyperactivity and anxiety-like behaviors of *Stxbp1^tm1d/+^*and *Stxbp1^tm1a/+^* mice in the open-field test. *Stxbp1^tm1d/+^*and *Stxbp1^tm1a/+^* mice showed an increase in the total moving distance (G) and speed (H), and a decrease in the ratio of center moving distances over total moving distance (I) and vertical activity (J). (**K**–**M**) *Stxbp1^tm1d/+^* and *Stxbp1^tm1a/+^* mice had weaker forelimb grip strength (K), made more foot slips per travel distance on a wire grid (L), and took more time to get down from a vertical pole (M). The numbers and ages of tested mice are indicated in the figures. Each filled (male) or open (female) circle represents one mouse. Bar graphs are mean ± s.e.m. * *P* < 0.05, ** *P* < 0.01, *** *P* < 0.001, **** *P* < 0.0001.

Homozygous mutants (*Stxbp1^tm1d/tm1d^*and *Stxbp1^tm1a/tm1a^*) died immediately after birth because they were completely paralyzed and could not breathe, consistent with the previous *Stxbp1* null alleles (Miyamoto et al., 2017; Verhage et al., 2000). Western blots with antibodies recognizing either the N- or C-terminus of Stxbp1 showed that Stxbp1 protein was absent in *Stxbp1^tm1d/tm1d^* and *Stxbp1^tm1a/tm1a^* mice at embryonic day 17.5 (***Figure 1-supplement 1B,C***), indicating that both alleles are null alleles. Importantly, both *Stxbp1^tm1d/+^*and *Stxbp1^tm1a/+^* mice showed a 50% reduction in Stxbp1 protein levels as compared to their wild type (WT) littermates at embryonic day 17.5 and 3 months of age (***Figure 1B*** and ***Figure 1-supplement 1B,C***), demonstrating that they are *Stxbp1* haploinsufficient mice. In theory, the KO and KO-first alleles could produce a truncated Stxbp1 protein of 18 kD and 16 kD, respectively. However, no such truncated proteins were observed in either heterozygous or homozygous mutants (***Figure 1-supplement 1B***), most likely because the truncated *Stxbp1* transcripts were degraded due to nonsense-mediated decay (Chang et al., 2007).

### *Stxbp1* haploinsufficient mice show a reduction in survival and body weights, and developed dystonia

We bred *Stxbp1^tm1d/+^* and *Stxbp1^tm1a/+^* mice with WT mice and found that at the time of genotyping (i.e., around postnatal week 3) *Stxbp1^tm1d/+^* and *Stxbp1^tm1a/+^* mice are 40% and 43% of the total offspring, respectively (***Figure 1-supplemtent 1D, Figure 1-supplemtent 2***), indicating a postnatal lethality phenotype. However, the lifespans of many mutant mice that survived through weaning were similar to those of WT littermates (***Figure 1-supplemtent 1E***). Thus, *Stxbp1* haploinsufficient mice show reduced survival, but this phenotype is not fully penetrant. *Stxbp1^tm1d/+^* and *Stxbp1^tm1a/+^* mice appeared smaller and their body weights were consistently about 20% less than their sex- and age-matched WT littermates (***Figure 1C,D***). At 4 weeks of age, *Stxbp1^tm1d/+^* and *Stxbp1^tm1a/+^* mice began to exhibit abnormal hindlimb clasping, indicative of dystonia. By 3 months of age, almost all mutant mice developed dystonia (***Figure 1C,E***). Thus, these observations indicate neurological deficits in *Stxbp1* haploinsufficient mice.

Guided by the symptoms of *STXBP1* encephalopathy human patients, we sought to perform behavioral and physiological assays to further examine the neurological and neuropsychiatric functions in male and female *Stxbp1* haploinsufficient mice. *Stxbp1^tm1d/+^* and *Stxbp1^tm1a/+^* mice were compared to their sex- and age-matched WT littermates.

### Impaired motor and normal sensory functions in *Stxbp1* haploinsufficient mice

Motor impairments including dystonia, ataxia, hypotonia, and tremor are frequently observed in *STXBP1* encephalopathy patients, but have not been recapitulated by the previous *Stxbp1* heterozygous knockout mice. Thus, we first assessed general locomotion by the open-field test where a mouse is allowed to freely explore an arena (***Figure 1F***). The locomotion of *Stxbp1^tm1d/+^* and *Stxbp1^tm1a/+^* mice was largely normal, but they traveled longer distances and faster than WT mice, indicating that *Stxbp1* haploinsufficient mice are hyperactive (***Figure 1G,H***). Both *Stxbp1^tm1d/+^* and *Stxbp1^tm1a/+^* mice explored the center region of the arena less than WT mice (***Figure 1I***) and made less vertical movements (***Figure 1J***), indicating that the mutant mice are more anxious. This anxiety phenotype was later confirmed by two other assays that specifically assess anxiety (see below). We used a variety of assays to further evaluate motor functions. *Stxbp1* haploinsufficient mice performed similarly to WT mice in the rotarod test, dowel test, inverted screen test, and wire hang test (***Figure 1-supplemtent 3***). However, the forelimb grip strength of *Stxbp1* haploinsufficient mice was weaker (***Figure 1K***). Furthermore, in the foot slip test where a mouse is allowed to walk on a wire grid, both *Stxbp1^tm1d/+^* and *Stxbp1^tm1a/+^* mice were not able to place their paws precisely on the wire to hold themselves and made many more foot slips than WT mice (***Figure 1L***). To assess the agility of mice, we performed the vertical pole test, which is often used to measure the bradykinesia of Parkinsonism. When mice were placed head-up on the top of a vertical pole, it took mutant mice much longer to orient themselves downward and descend the pole than WT mice (***Figure 1M***). Together, these results indicate that *Stxbp1* haploinsufficient mice do not develop ataxia, but their fine motor coordination and muscle strength are reduced.

We next examined the acoustic sensory function and found that *Stxbp1^tm1d/+^* and *Stxbp1^tm1a/+^* mice showed normal startle responses to different levels of sound (***Figure 1-supplemtent 4A***). To test sensorimotor gating, we measured the pre-pulse inhibition where the startle response to a strong sound is reduced by a preceding weaker sound. *Stxbp1^tm1d/+^*and *Stxbp1^tm1a/+^* mice displayed similar pre-pulse inhibition as WT mice (***Figure 1-supplemtent 4B***). They also had normal nociception as measured by the hot plate test (***Figure 1-supplemtent 4C***). Thus, the sensory functions and sensorimotor gating of *Stxbp1* haploinsufficient mice are normal.

### Cognitive functions of *Stxbp1* haploinsufficient mice are severely impaired

Intellectual disability is a core feature of *STXBP1* encephalopathy, as all patients are intellectually disabled and the vast majority are severe to profound (Stamberger et al., 2016). However, the learning and memory deficits described in the previous *Stxbp1* heterozygous knockout mice are mild and inconsistent (Kovačević et al., 2018; Miyamoto et al., 2017; Orock et al., 2018). To assess cognitive functions, we tested *Stxbp1* haploinsufficient mice in three different paradigms, object recognition, associative learning and memory, and working memory. First, we performed the novel object recognition test that exploits the natural tendency of mice to explore novel objects to evaluate their memories. This task is thought to depend on the hippocampus and cortex (Antunes and Biala, 2012; Cohen and Stackman, 2015). When tested with an inter-trial interval of 24 hours, WT mice interacted more with the novel object than the familiar object, whereas *Stxbp1^tm1d/+^*and *Stxbp1^tm1a/+^* mice interacted equally between the familiar and novel objects (***Figure 2A***). We also evaluated *Stxbp1^tm1d/+^* mice with an inter-trial interval of 5 minutes and observed a similar deficit (***Figure 2-supplement 1A***). We noticed that mutant mice overall spent less time interacting with the objects than WT mice during the trials (***Figure 2-supplement 1B***), which might reduce their “memory load” of the objects. We hence allowed *Stxbp1^tm1d/+^* mice to spend twice as much time as WT mice in each trial to increase their interaction time with the objects, but they still showed a similar deficit in recognition memory (***Figure 2-supplement 1D***). Thus, both long-term and short-term recognition memories are impaired in *Stxbp1* haploinsufficient mice.

**Figure 2.**
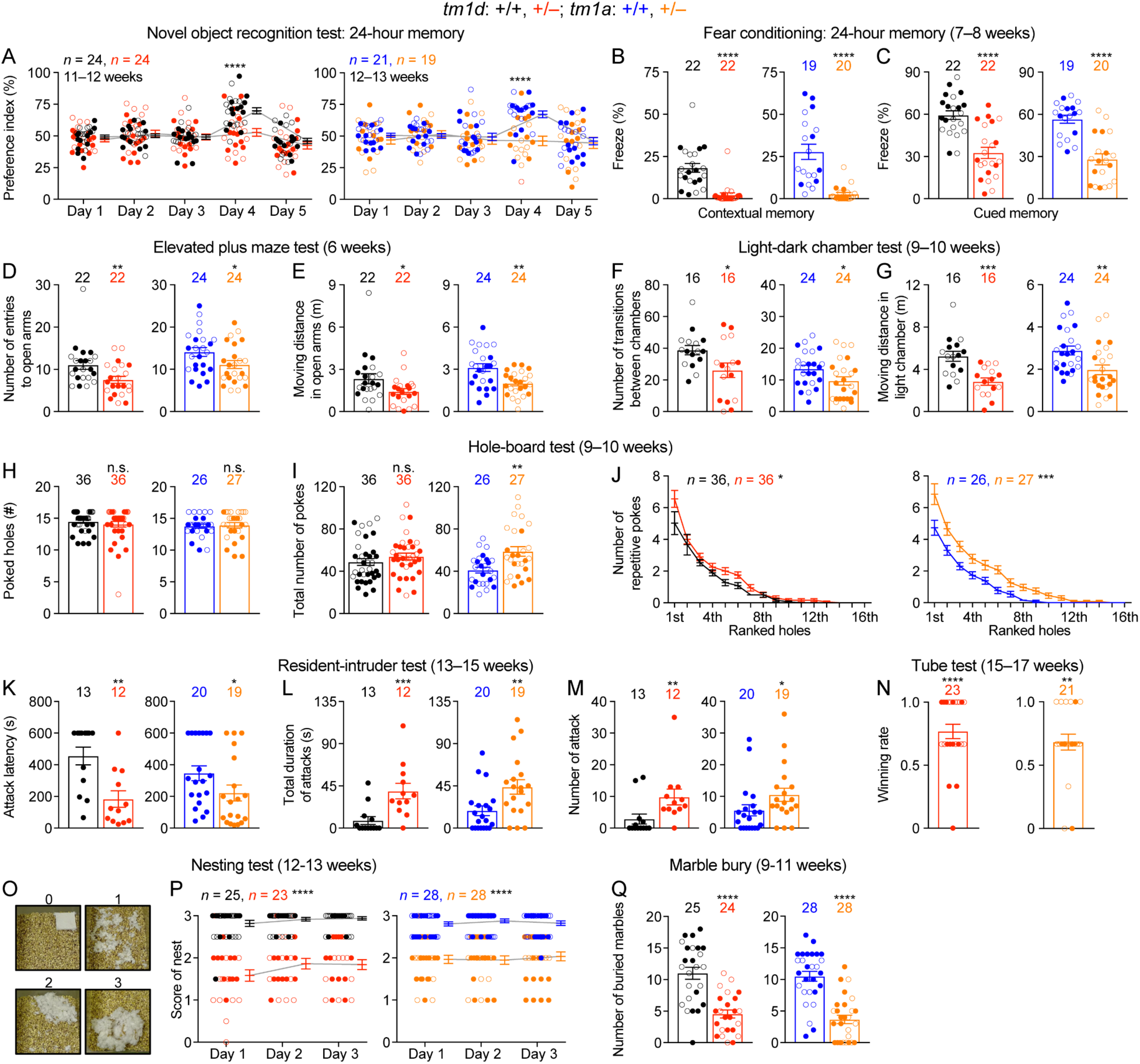
*Stxbp1* haploinsufficient mice show impaired cognition, increased anxiety-like, repetitive, and aggressive behaviors, and reduced nest building and digging behaviors. (**A**) In the novel object recognition test with 24-hour testing intervals, in contrast to WT mice, *Stxbp1^tm1d/+^*and *Stxbp1^tm1a/+^* mice did not show a preference for the novel object on day 4 when they were presented with a familiar and a novel object. (**B,C**) In the fear conditioning test, *Stxbp1^tm1d/+^*and *Stxbp1^tm1a/+^* mice showed a reduction in both context- and cue-induced freeze 24 hours after training. (**D,E**) In the elevated plus maze test, *Stxbp1^tm1d/+^*and *Stxbp1^tm1a/+^* mice entered the open arms less frequently (D) and traveled shorter distance in the open arms (E). (**F,G**) In the light-dark chamber test, *Stxbp1^tm1d/+^* and *Stxbp1^tm1a/+^* mice made less transitions between the light and dark chambers (F) and traveled shorter distance in the light chamber (G). (**H–J**) In the hole-board test, *Stxbp1^tm1d/+^* and *Stxbp1^tm1a/+^* mice poked similar numbers of holes as WT mice (H) and made similar or more total nose pokes (I). They made more repetitive nose pokes (i.e., ≥ 2 consecutive pokes) than WT mice across different holes (J). (**K–M**) In the resident-intruder test, male *Stxbp1^tm1d/+^* and *Stxbp1^tm1a/+^* mice showed a reduction in the latency to attack the male intruder mice (K). The total duration (L) and number (M) of their attacks were increased as compared to WT mice. (**N**) In the tube test, *Stxbp1^tm1d/+^* and *Stxbp1^tm1a/+^* mice won more competitions against their WT littermates. (**O,P**) *Stxbp1^tm1d/+^* and *Stxbp1^tm1a/+^* mice built poor quality nests. The quality of the nests was scored according to the criteria in (O) for 3 consecutive days (P). (**Q**) *Stxbp1^tm1d/+^*and *Stxbp1^tm1a/+^* mice buried fewer marbles than WT mice. The numbers and ages of tested mice are indicated in the figures. Each filled (male) or open (female) circle represents one mouse. Bar graphs are mean ± s.e.m. n.s. *P* > 0.05, * *P* < 0.05, ** *P* < 0.01, *** *P* < 0.001, **** *P* < 0.0001.

Second, we used the Pavlovian fear conditioning paradigm to evaluate associative learning and memory, in which a mouse learns to associate a specific environment (i.e., the context) and a sound (i.e., the cue) with electric foot shocks. The fear memory is manifested by the mouse freezing when it is subsequently exposed to this specific context or cue without electric shocks. At two tested ages, *Stxbp1^tm1d/+^* and *Stxbp1^tm1a/+^* mice displayed a profound reduction in both context- and cue-induced freeze when tested 24 hours after the conditioning (***Figure 2B,C, Figure 2-supplement 1E,F***). We also tested *Stxbp1^tm1d/+^* mice 1 hour after the conditioning and observed similar deficits (***Figure 2-supplement 1G***). Since the acoustic startle response and nociception are intact in *Stxbp1* haploinsufficient mice (***Figure 1-supplemtent 4C***), these results indicate that *Stxbp1* haploinsufficiency impairs both hippocampus-dependent contextual and hippocampus-independent cued fear memories.

Finally, we used the Y maze spontaneous alternation test to examine working memory, but did not observe significant difference between *Stxbp1^tm1d/+^* and WT mice (***Figure 2-supplement 1H***). Taken together, our results indicate that both long-term and short-term forms of recognition and associative memories are severely impaired in *Stxbp1* haploinsufficiency mice, but their working memory is intact.

### *Stxbp1* haploinsufficient mice exhibit an increase in anxiety-like and repetitive behaviors

A number of neuropsychiatric phenotypes including hyperactivity, anxiety, stereotypies, aggression, and autistic features were reported in subsets of *STXBP1* encephalopathy patients. We used a battery of behavioral assays to characterize each of these features in *Stxbp1* haploinsufficiency mice. The open-field test indicates that *Stxbp1* haploinsufficiency mice are hyperactive and more anxious than WT mice (***Figure 1F–J***). To specifically assess anxiety-like behaviors, we tested *Stxbp1^tm1d/+^*and *Stxbp1^tm1a/+^* mice in the elevated plus maze and light-dark chamber tests where a mouse is allowed to explore the open or closed arms of the maze and the clear or black chamber of the box, respectively. *Stxbp1^tm1d/+^*and *Stxbp1^tm1a/+^* mice entered the open arms and clear chamber less frequently and and traveled shorter distance in the open arms and clear chamber than WT mice (***Figure 2D–G; Figure 2-supplement 1I,J***). Hence, these results confirm the heightened anxiety in *Stxbp1* haploinsufficient mice and are consistent with the previous studies (Hager et al., 2014; Kovačević et al., 2018; Miyamoto et al., 2017).

To assess the stereotypy and repetitive behaviors, we used the hole-board test to measures the pattern of mouse exploratory nose poke (also called head dipping) behavior. As compared to WT mice, *Stxbp1* haploinsufficient mice explored similar numbers of holes (***Figure 2H***) and made similar or larger numbers of nose pokes (***Figure 2I***). We analyzed the repetitive nose pokes (i.e., ≥ 2 consecutive pokes) into the same hole as a measure of repetitive behaviors. The mutant mice made more repetitive nose pokes than WT mice across many holes (***Figure 2J***), indicating that *Stxbp1* haploinsufficiency in mice causes abnormal stereotypy and repetitive behaviors, a neuropsychiatric feature observed in about 20% of the *STXBP1* encephalopathy patients (Stamberger et al., 2016).

### Social aggression of *Stxbp1* haploinsufficient mice are elevated

During daily mouse husbandry, we noticed incidences of fighting and injuries of WT and *Stxbp1* haploinsufficient mice in their home cages when *Stxbp1* haploinsufficient mice were present. No injuries were observed when *Stxbp1* haploinsufficient mice were singly housed, suggesting that the injuries likely resulted from fighting instead of self-injury. To formally examine aggressive behaviors, we first performed the resident-intruder test, in which a male intruder mouse is introduced into the home cage of a male resident mouse, and the aggressive behaviors of the resident towards the intruder were scored. As compared to WT mice, male resident *Stxbp1^tm1d/+^* and *Stxbp1^tm1a/+^* mice were more likely to attack and spent more time attacking the intruders (***Figure 2K–M***). Another paradigm to assess aggression and social dominance is the tube test, in which two mice are released into the opposite ends of a tube, and the more dominant and aggressive mouse will win the competition by pushing its opponent out of the tube. When *Stxbp1^tm1d/+^* and *Stxbp1^tm1a/+^* mice were placed against their sex- and age-matched WT littermates, *Stxbp1* haploinsufficient mice won more competitions despite their smaller body sizes (***Figure 2N***). Thus, *Stxbp1* haploinsufficiency elevates innate aggression in mice.

To further evaluate social interaction, we performed the three-chamber test where a mouse is allowed to interact with an object or a sex- and age-matched partner mouse. Like WT mice, *Stxbp1^tm1d/+^*and *Stxbp1^tm1a/+^* mice preferred to interact with the partner mice rather than the objects (***Figure 2-supplement 1K***), indicating that *Stxbp1* haploinsufficiency does not compromise general sociability. Interestingly, the mutant mice in fact spent significantly more time than WT mice interacting with the partner mice (*P* < 0.0001 for *Stxbp1^tm1d/+^* vs. WT and *P* = 0.0015 for *Stxbp1^tm1a/+^* vs. WT), which might be due to the increased aggression of the mutant mice. Furthermore, we used the partition test to examine the preference for social novelty, in which a mouse is allowed to interact with a familiar or novel partner mouse. Both WT and *Stxbp1^tm1d/+^* mice preferentially interacted more with the novel partner mice (***Figure 2-supplement 1L***). These results indicate that the general sociability and interest in social novelty are normal in *Stxbp1* haploinsufficient mice.

### Reduced nest building and digging behaviors in *Stxbp1* haploinsufficient mice

To further assess the well-being and neuropsychiatric phenotypes of *Stxbp1* haploinsufficient mice, we performed the Nestlet shredding test and marble burying test to examine two innate behaviors, nest building and digging, respectively. We provided a Nestlet (pressed cotton square) to each mouse in the home cage and scored the degree of shredding and nest quality after 24, 48, and 72 hours (***Figure 2O***). *Stxbp1^tm1d/+^*and *Stxbp1^tm1a/+^* mice consistently scored lower than WT mice at all time points (***Figure 2P***). In the marble burying test, the *Stxbp1^tm1d/+^* and *Stxbp1^tm1a/+^* mice buried fewer marbles than WT mice (***Figure 2Q***). The interpretation of marble burying remains controversial, as it may measure anxiety, compulsive-like behavior, or simply digging behavior (Deacon, 2006; Thomas et al., 2009; Wolmarans et al., 2016). Since *Stxbp1* haploinsufficient mice show elevated anxiety and repetitive behaviors, the reduced marble burying likely reflects an impairment of digging behavior, possibly due to the motor deficits. Likewise, the motor deficits may also contribute to the reduced nest building behavior.

### Cortical hyperexcitability and epileptic seizures in *Stxbp1* haploinsufficient mice

Another core feature of *STXBP1* encephalopathy is epilepsy with a broad spectrum of seizure types, such as epileptic spasm, focal, tonic, clonic, myoclonic, and absence seizures (Stamberger et al., 2016; Suri et al., 2017). To investigate if *Stxbp1* haploinsufficient mice have abnormal cortical activity and epileptic seizures, we performed chronic video-electroencephalography (EEG) and electromyography (EMG) recordings in freely moving *Stxbp1^tm1d/+^* mice and their sex- and age-matched WT littermates. We implanted three EEG electrodes in the frontal and somatosensory cortices and an EMG electrode in the neck muscle to record intracranial EEG and EMG, respectively, for at least 72 hours (***Figure 3A***). The phenotypes of each mouse are summarized in ***Supplementary Table 1***. *Stxbp1^tm1d/+^* mice exhibited cortical hyperexcitability and several epileptiform activities. First, they had numerous spike-wave discharges (SWDs) that typically were 3–6 Hz and lasted 1–2 s (***Figure 3C,E,F***). These oscillations showed similar characteristics to those generalized spike-wave discharges observed in animal models of absence seizures (Depaulis and Charpier, 2018; Maheshwari and Noebels, 2014). A much smaller number of SWDs with similar characteristics were also observed in WT mice (***Figure 3B***), consistent with previous studies (Arain et al., 2012; Letts et al., 2014). On average, the frequency of SWD episodes in *Stxbp1^tm1d/+^* mice was more than 40 folds of that in WT mice (***Figure 3E,F***). Importantly, SWDs frequently occurred in a cluster manner (i.e., ≥ 5 episodes with an inter-episode-interval of ≤ 60 s) in *Stxbp1^tm1d/+^* mice, which never occurred in WT mice (***Figure 3-supplement 1; Figure 3-supplement 2 Video S1***). Furthermore, 56 episodes of SWDs from 10 out of 13 *Stxbp1^tm1d/+^* mice lasted more than 4 s, among which 54 episodes occurred during rapid eye movement (REM) sleep (***Figure 3D; Figure 3-supplement 3 Video S2***) and the other 2 episodes occurred when mice were awake. In contrast, only 1 out of 11 WT mice had 3 episodes of such long SWDs, all of which occurred when mice were awake (***Supplementary Table 1***). In *Stxbp1^tm1d/+^* mice, SWDs were most frequent during the nights, but occurred throughout the days and nights (***Figure 3F***), indicating a general cortical hyperexcitability and abnormal synchrony in *Stxbp1* haploinsufficient mice.

**Figure 3.**
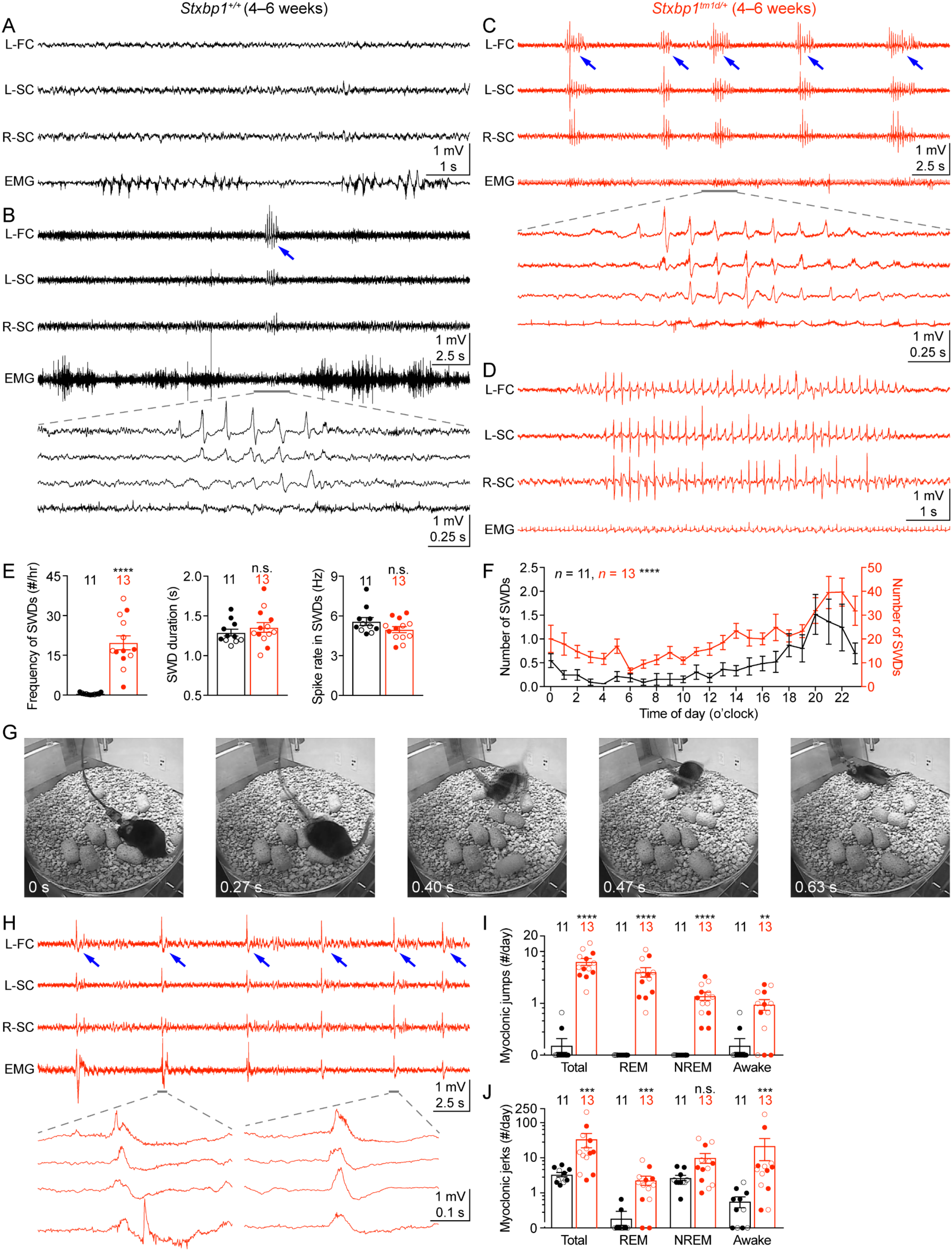
*Stxbp1^tm1d/+^* mice exhibit cortical hyperexcitability and epileptic seizures. (**A–D**) Representative EEG traces of the left frontal cortex (L-FC), left somatosensory cortex (L-SC), and right somatosensory cortex (R-SC), and EMG traces from WT (A,B) and *Stxbp1^tm1d/+^* mice (C,D). Spike-wave discharges (SWDs, indicated by the blue arrows) occurred frequently and often in a cluster manner in *Stxbp1^tm1d/+^* mice (see ***Figure 3-supplement 2 Video S1***). The grey line-highlighted SWDs from WT and *Stxbp1^tm1d/+^* mice were expanded to show the details of the oscillations (B,C). A long SWD (i.e., > 4 s) during REM sleep from a *Stxbp1^tm1d/+^* mouse is shown in (D) (see ***Figure 3-supplement 3 Video S2***). (**E**) Summary data showing the overall SWD frequency (left panel), duration (middle panel), and average spike rate (right panel). (F) The numbers of SWDs per hour in WT (left Y axis) and *Stxbp1^tm1d/+^* (right Y axis) mice are plotted as a function of time of day and averaged over 3 days. (**G**) Video frames showing a myoclonic jump from a *Stxbp1^tm1d/+^* mouse (see ***Figure 3-supplement 4 Video S3***). The mouse was in REM sleep before the jump. (**H**) Representative EEG and EMG traces showing myoclonic jerks (indicated by the blue arrows) from a *Stxbp1^tm1d/+^* mouse (see ***Figure 3-supplement 5 Video S4***). Two episodes of myoclonic jerks highlighted by the grey lines were expanded to show that the EEG discharges occurred prior to (the first episode) or simultaneously with (the second episode) the EMG discharges. (**I,J**) Summary data showing the frequencies of two types of myoclonic seizures in different behavioral states. The numbers and ages of recorded mice are indicated in the figures. Each filled (male) or open (female) circle represents one mouse. Bar graphs are mean ± s.e.m. n.s. *P* > 0.05, ** *P* < 0.01, *** *P* < 0.001, **** *P* < 0.0001.

Second, *Stxbp1^tm1d/+^* mice experienced frequent myoclonic seizures that were manifested as sudden jumps or more subtle, involuntary muscle jerks associated with EEG discharges (***Figure 3G,H***). The large movement artifacts associated with the myoclonic jumps precluded proper interpretation of EEG signals, but this type of myoclonic seizures was observed in all 13 recorded *Stxbp1^tm1d/+^* mice and the majority of episodes occurred during REM or non-rapid eye movement (NREM) sleep (***Figure 3I; Figure 3-supplement 4 Video S3***). There were 3 similar jumps in 2 out of 11 WT mice that were indistinguishable from those in *Stxbp1^tm1d/+^* mice, but all of them occurred when mice were awake (***Figure 3I***). Moreover, the more subtle myoclonic jerks occurred frequently and often in clusters in *Stxbp1^tm1d/+^* mice, whereas only isolated events were observed in WT mice at a much lower frequency (***Figure 3H,J; Figure 3-supplement 5 Video S4***). EEG and EMG recordings showed that the cortical EEG spikes associated with the myoclonic jerks occurred before or simultaneously with the neck muscle EMG discharges (***Figure 3H***), consistent with the cortical or subcortical origins of myoclonuses, respectively (Avanzini et al., 2016).

### *Stxbp1* haploinsufficiency reduces synaptic inhibition in a cell-type specific manner

To identify cellular mechanisms that may underlie the cortical hyperexcitability and neurological deficits in *Stxbp1* haploinsufficient mice, we examined neuronal excitability and synaptic transmission in the somatosensory cortex. Whole-cell current clamp recordings of layer 2/3 pyramidal neurons in acute brain slices revealed only a small increase in the input resistances of *Stxbp1^tm1d/+^* neurons as compared to WT neurons (***Figure 4-supplement 1***). Previous studies showed that synaptic transmission was reduced in the cultured hippocampal neurons from heterozygous *Stxbp1* knockout mice and human neurons derived from heterozygous *STXBP1* knockout embryonic stem cells (Orock et al., 2018; Patzke et al., 2015; Toonen et al., 2006). However, such a decrease in excitatory transmission is unlikely adequate to explain how *Stxbp1* haploinsufficiency *in vivo* leads to cortical hyperexcitability. Thus, we focused on the inhibitory synaptic transmission originating from two major classes of cortical inhibitory neurons, Pv and Sst interneurons. A Cre-dependent tdTomato reporter line, *Rosa26-CAG-LSL-tdTomato* (Madisen et al., 2010), and *Pv-ires-Cre* (Hippenmeyer et al., 2005) or *Sst-ires-Cre* (Taniguchi et al., 2011) were used to identify Pv or Sst interneurons, respectively. We used whole-cell current clamp to stimulate a single Pv or Sst interneuron in layer 2/3 with a brief train of action potentials and whole-cell voltage clamp to record the resulting unitary inhibitory postsynaptic currents (uIPSCs) in a nearby pyramidal neuron (***Figure 4A,E***). The connectivity rate of Pv interneurons to pyramidal neurons was unaltered in *Stxbp1^tm1d/+^;Rosa26^tdTomato/+^;Pv^Cre/+^* mice (***Figure 4B***), but the unitary connection strength was reduced by 45% as compared to *Stxbp1^+/+^;Rosa26^tdTomato/+^;Pv^Cre/+^* mice (***Figure 4C***). In contrast, *Stxbp1^tm1d/+^;Rosa26^tdTomato/+^;Sst^Cre/+^* mice showed a 26% reduction in the connectivity rate of Sst interneurons to pyramidal neurons (***Figure 4F***), but the unitary connection strength was normal (***Figure 4G***). The short-term synaptic depression of both inhibitory connections during the train of stimulations was normal (***Figure 4D,H***). Thus, cortical inhibition mediated by Pv and Sst interneurons is impaired in *Stxbp1* haploinsufficient mice, representing a likely cellular mechanism for the cortical hyperexcitability and neurological deficits.

**Figure 4.**
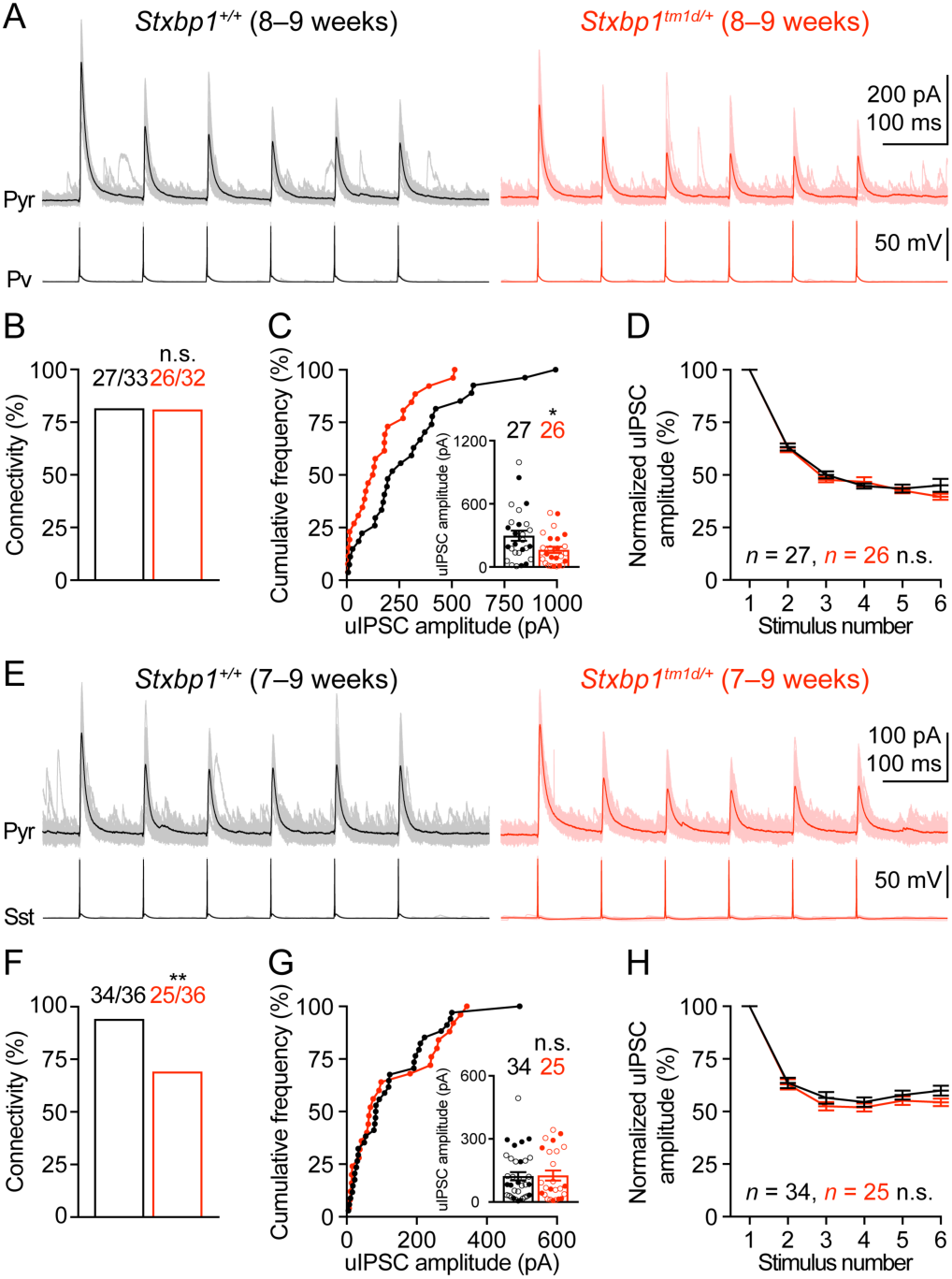
Inhibitory synapses from Pv and Sst interneurons are differentially impaired in *Stxbp1^tm1d/+^* mice. (**A**) uIPSCs of a layer 2/3 pyramidal neuron in the somatosensory cortex (upper panels) evoked by a train of 10-Hz action potentials in a nearby Pv interneuron (lower panels) from WT and *Stxbp1^tm1d/+^* mice. 50 individual traces (lighter color) and the average trace (darker color) are superimposed. Note smaller uIPSCs in the *Stxbp1^tm1d/+^* neuron. (**B**) Unitary connectivity rates from Pv interneurons to pyramidal neurons were similar between WT (27 connections out of 33 pairs) and *Stxbp1^tm1d/+^* (26 connections out of 32 pairs) mice. (**C**) Cumulative frequencies of uIPSC amplitudes evoked by the first action potentials in the trains (median: WT, 217.3 pA; *Stxbp1^tm1d/+^*, 127.1 pA). Inset, each filled (male) or open (female) circle represents the uIPSC amplitude of one synaptic connection. (**D**) uIPSC amplitudes during the trains of action potentials were normalized by the amplitudes of the first uIPSCs. Note the similar synaptic depression between WT and *Stxbp1^tm1d/+^* neurons. (**E–H**) Similar to (A–D), but for Sst interneurons. Unitary connectivity rates from Sst interneurons to pyramidal neurons (F) in *Stxbp1^tm1d/+^* mice (25 connections out of 36 pairs) were less than WT mice (34 connections out of 36 pairs). The uIPSC amplitudes evoked by the first action potentials in the trains (G, median: 83.5 pA and 68.0 pA, respectively) and synaptic depression (H) were similar between WT and *Stxbp1^tm1d/+^* mice. Bar graphs are mean ± s.e.m. n.s. *P* > 0.05, * *P* < 0.05, ** *P* < 0.01.

## Discussion

Extensive biochemical and structural studies of Stxbp1/Munc18-1 have elucidated its crucial role in synaptic vesicle exocytosis (Rizo and Xu, 2015), but provided little insight into its functional role at the organism level. Hence, apart from being an essential gene, the significance of *STXBP1* dysfunction *in vivo* was not appreciated until its *de novo* heterozygous mutations were discovered first in epileptic encephalopathies (Saitsu et al., 2008) and later in other neurodevelopmental disorders (Deciphering Developmental Disorders Study, 2015; Hamdan et al., 2011; 2009; Rauch et al., 2012). In this study, we generated two lines of *Stxbp1* haploinsufficient mice (*Stxbp1^tm1d/+^* and *Stxbp1^tm1a/+^*) and systematically characterized them in all of the neurological and neuropsychiatric domains affected by *STXBP1* encephalopathy. These mice exhibit reduced survival, hindlimb clasping, impaired motor coordination, learning and memory deficits, hyperactivity, increased anxiety-like and repetitive behaviors, aggression, and epileptic seizures. Sensory abnormality has not been documented in *STXBP1* encephalopathy patients (Stamberger et al., 2016) and we also did not observe any sensory dysfunctions in *Stxbp1* haploinsufficient mice. Thus, despite the large phenotypic spectrum of *STXBP1* encephalopathy in humans, our *Stxbp1* haploinsufficient mice recapitulate all key features of this neurodevelopmental disorder and are construct and face valid models of *STXBP1* encephalopathy. About 20% of the *STXBP1* encephalopathy patients showed autistic traits (Stamberger et al., 2016), but we and others (Kovačević et al., 2018; Miyamoto et al., 2017) did not observe an impairment of social interaction in mutant mice using the three-chamber and partition tests. Perhaps the elevated aggression in *Stxbp1* haploinsufficient mice confounds these tests, or new mouse models that more precisely mimic the genetic alterations in that subset of *STXBP1* encephalopathy patients are required to recapitulate this phenotype.

Prior studies using the other three lines of *Stxbp1* heterozygous knockout mouse models reported only a subset of the neurological and neuropsychiatric deficits that we observed here (Hager et al., 2014; Kovačević et al., 2018; Miyamoto et al., 2017; Orock et al., 2018). For example, the reduced survival, hindlimb clasping, motor dysfunction, and increased repetitive behavior were not documented in the previous models. The previously reported cognitive phenotypes were much milder than what we observed. Both *Stxbp1^tm1d/+^* and *Stxbp1^tm1a/+^* mice showed severe impairments in the novel objection recognition and fear conditioning tests. In contrast, another line of *Stxbp1* heterozygous knockout mice showed normal spatial learning in the Morris water maze and Barnes maze (a dry version of the spatial maze) in one study (Kovačević et al., 2018), but reduced spatial learning and memory in the radial arm water maze in another study (Orock et al., 2018). Different behavioral tests could have contributed to such differences among studies. However, a subtle but perhaps key difference is the Stxbp1 protein levels in different lines of heterozygous mutant mice. Stxbp1 is reduced by 50% in both of our *Stxbp1^tm1d/+^*and *Stxbp1^tm1a/+^* mice, but only by 25–40% in other heterozygous knockout mice (Miyamoto et al., 2017; Orock et al., 2018), which may lead to fewer or less severe phenotypes in the previous models.

Dysfunction of cortical GABAergic inhibition has been widely considered as a primary defect in animal models of autism spectrum disorder, schizophrenia, Down syndrome, and epilepsy among other neurological disorders (Contestabile et al., 2017; Lee et al., 2017; Marín, 2012; Nelson and Valakh, 2015; Paz and Huguenard, 2015; Ramamoorthi and Lin, 2011). In many cases, the origins of GABAergic dysfunction were either unidentified or attributed to Pv interneurons. Sst interneurons have only been directly implicated in a few disease models (Ito-Ishida et al., 2015; Rubinstein et al., 2015) despite their important physiological functions. Here we identified distinct deficits at Pv and Sst interneuron synapses in *Stxbp1* haploinsufficient mice, suggesting that Stxbp1 may have diverse functions at distinct synapses. The reduction in the strength of Pv interneuron synapses is consistent with the previous results that basal synaptic transmission is reduced at the neuromuscular junctions of *Stxbp1* heterozygous null flies and mice (Toonen et al., 2006; Wu et al., 1998) and the glutamatergic synapses of human *STXBP1* heterozygous knockout neurons (Patzke et al., 2015). The reduced synaptic strength is likely due to a decrease in the number of readily releasable vesicles or release probability given the crucial role of Stxbp1 in synaptic vesicle priming and fusion (Rizo and Xu, 2015). On the other hand, the reduction in the connectivity of Sst interneuron synapses is unexpected, as Stxbp1 has not yet been implicated in the formation or maintenance of synapses. Complete loss of Stxbp1 in mice does not appear to affect the initial formation of neural circuits, but causes cell-autonomous neurodegeneration and protein trafficking defects (Law et al., 2016; Verhage et al., 2000). Since Munc13-1/2 double knockout mice also lack synaptic exocytosis, but do not show neurodegeneration (Varoqueaux et al., 2002), the degeneration phenotype in Stxbp1 null mice is unlikely caused by the total arrest of synaptic exocytosis. Thus, Stxbp1 may regulate other intracellular processes in addition to presynaptic transmitter release, and we speculate that it may be involved in a protein trafficking process important for the formation or maintenance of Sst interneuron synapses. Nevertheless, the impairment of Pv and Sst interneuron-mediated inhibition likely constitutes a key mechanism underlying the cortical hyperexcitability and neurobehavioral phenotypes of *Stxbp1* haploinsufficient mice. Future studies using cell-type specific *Stxbp1* haploinsufficient mouse models will help determine the role of GABAergic interneurons in the disease pathogenesis.

There are over 100 developmental brain disorders that arise from mutations in postsynaptic proteins, whereas mutations in much fewer presynaptic proteins have been identified to cause neurodevelopmental disorders (Bayés et al., 2011; Deciphering Developmental Disorders Study, 2017). However, in addition to STXBP1, pathogenic variants in other key components of the presynaptic neurotransmitter release machinery were recently discovered in neurodevelopmental disorders. These include Ca^2+^-sensor synaptotagmin 1 (SYT1), vesicle priming factor unc-13 homolog A (UNC13A), and all three components of the neuronal SNAREs, syntaxin 1B (STX1B), synaptosome associated protein 25 (SNAP25), and vesicle associated membrane protein 2 (VAMP2) (Baker et al., 2015; 2018; Engel et al., 2016; Fukuda et al., 2018; Hamdan et al., 2017; Lipstein et al., 2017; Rohena et al., 2013; Salpietro et al., 2019; Schubert et al., 2014; Shen et al., 2014; Wolking et al., 2019). Haploinsufficiency of these synaptic proteins is likely the leading disease mechanism because the majority of the cases were caused by heterozygous loss-of-function mutations. The clinical features of these disorders are diverse, but significantly overlap with those of *STXBP1* encephalopathy. The most common phenotypes are intellectual disability and epilepsy or cortical hyperexcitability, which can be considered as the core features of these genetic synaptopathies. Thus, *Stxbp1* haploinsufficient mice are a valuable model to understand the cellular and circuit origins of these complex disorders and a growing list of neurodevelopmental disorders caused by synaptic dysfunction.

## Methods

### Mice

We obtained *Stxbp1^tm1a(EUCOMM)Hmgu^* embryonic stem (ES) cell clones (C57Bl/6N strain) from the European Conditional Mouse Mutagenesis Program (EUCOMM) and confirmed the targeting by Southern blots. Two ES cell clones (HEPD0510_5_A09 and HEPD0510_5_B10) were injected into blastocysts to generate chimeric mice. We obtained germline transmission from the clone HEPD0510_5_A09 by breeding the chimeric mice to B6(Cg)-Tyrc-2J/J mice (JAX #000058) and established the KO-first (*tm1a*) line. Heterozygous KO-first mice were crossed to *Rosa26-Flpo* mice (Raymond and Soriano, 2007) to remove the trapping cassette in the germline. The resulting offspring were then crossed to *Sox2-Cre* mice (Hayashi et al., 2002) to delete exon 7 in the germline to generate the KO (*tm1d*) line. Both *Rosa26-Flpo* and *Sox2-Cre* mice were obtained from the Jackson Laboratory (#012930 and 008454, respectively). *Stxbp1* mice were genotyped by PCR using primer sets 5’-TTCCACAGCCCTTTACAGAAAGG-3’ and 5’-ATGTGTATGCCTGGACTCACAGGG-3’ for WT allele, 5’-TTCCACAGCCCTTTACAGAAAGG-3’ and 5’-CAACGGGTTCTTCTGTTAGTCC-3’ for KO-first allele, and 5’-TTCCACAGCCCTTTACAGAAAGG-3’ and 5’-TGAACTGATGGCGAGCTCAGACC-3’ for KO allele.

Heterozygous *Stxbp1* KO-first and KO mice were crossed to wild type (WT) C57BL/6J mice (JAX #000664) for maintaining both lines on the C57BL/6J background and for generating experimental cohorts. Male BALB/cAnNTac mice were obtained from Taconic (#BALB-M). *Pv-ires-Cre* (Hippenmeyer et al., 2005), *Sst-ires-Cre* (Taniguchi et al., 2011), and *Rosa26-CAG-LSL-tdTomato* (Madisen et al., 2010) mice were obtained from the Jackson Laboratory (#017320, 013044, and 007914, respectively). *Pv-ires-Cre* and *Rosa26-CAG-LSL-tdTomato* mice were maintained on the C57BL/6J background. *Sst-ires-Cre* mice were on a C57BL/6;129S4 background. Heterozygous KO mice were crossed to *Rosa26-CAG-LSL-tdTomato* mice to generate *Stxbp1^tm1d/+^;Rosa26^tdTomato/tdTomato^* mice. *Pv-ires-Cre* and *Sst-ires-Cre* mice were then crossed to *Stxbp1^tm1d/+^;Rosa26^tdTomato/tdTomato^* mice to generate *_Stxbp1tm1d/+;Rosa26tdTomato/+;PvCre/+_* _or *Stxbp1*_*_+/+;Rosa26tdTomato/+;PvCre/+_* _and_ *Stxbp1^tm1d/+^;Rosa26^tdTomato/+^;Sst^Cre/+^*or *Stxbp1^+/+^;Rosa26^tdTomato/+^;Sst^Cre/+^* mice, respectively.

Mice were housed in an Association for Assessment and Accreditation of Laboratory Animal Care International-certified animal facility on a 14-hour/10-hour light/dark cycle. All procedures to maintain and use mice were approved by the Institutional Animal Care and Use Committee at Baylor College of Medicine.

### Southern and Western blots

Southern and Western blot analyses were performed according standard protocols. For Southern blots, genomic DNA was extracted from ES cells and digested with BspHI for the 5’ probe or MfeI for the 3’ probe (***Figure 1-supplement 1A***). ^32^P-labeled probes were used to detect DNA fragments. For Western blots, proteins were extracted from the brains at embryonic day 17.5 or 3 months of age using lysis buffer containing 50 mM Tris-HCl (pH 7.6), 150 mM NaCl, 1 mM EDTA, 1% Triton X-100, 0.5% Na-deoxycholate, 0.1% SDS, and 1 tablet of cOmplete™, Mini, EDTA-free Protease Inhibitor Cocktail (Roche) in 10 ml buffer. Stxbp1 was detected by a rabbit antibody against the N terminal residues 58–70 (Abcam, catalog # ab3451, lot #GR79394-18, 1:2,000 or 1:5,000 dilution) or a rabbit antibody against the C terminal residues 580–594 (Synaptic Systems, catalog # 116002, lot # 116002/15, 1:2,000 or 1:5,000 dilution). Gapdh was detected by a rabbit antibody (Santa Cruz Biotechnology, catalog # sc-25778, lot # A0515, 1:300 or 1:1,000 dilution). Primary antibodies were detected by a goat anti-rabbit antibody conjugated with IRDye 680LT (LI-COR Biosciences, catalog # 925-68021, lot # C40917-01, 1:20,000 dilution). Proteins were visualized and quantified using an Odyssey CLx Imager and Image Studio Lite version 5.0 (LI-COR Biosciences).

### Behavioral Tests

All behavioral experiments except the tube test were performed and analyzed blind to the genotypes. Approximately equal numbers of *Stxbp1* mutant mice and their sex- and age-matched WT littermates of both sexes were tested in parallel in each experiment except the resident-intruder test where only male mice were used. In each cage, two mutant and two WT mice were housed together. Before all behavioral tests, mice were habituated in the behavioral test facility for at least 30 minutes. Gender effect was inspected by two-way or three-way ANOVA. If both sexes showed similar phenotypes, the data were aggregated together to simplify the presentation; otherwise they were presented separately. The ages of the tested mice were indicated in figures.

#### Open-field test

A mouse was placed in the center of a clear, open chamber (40 × 40 × 30 cm) and allowed to freely explore for 30 minutes in the presence of 700–750-lux illumination and 65-dB background white noise. In each chamber, two layers of light beams (16 for each layer) in the horizontal X and Y directions capture the locomotor activity of the mouse. The horizontal plane was evenly divided into 256 squares (16 × 16), and the center zone is defined as the central 100 squares (10 × 10). The horizontal travel and vertical activity were quantified by either an Open Field Locomotor system or a VersaMax system (OmniTech).

#### Rotarod test

A mouse was placed on an accelerating rotarod apparatus (Ugo Basile). Each trial lasted for a maximum of 5 minutes, during which the rod accelerated linearly from 4 to 40 revolutions per minute (RPM) or 8 to 80 RPM. The time when the mouse walks on the rod and the latency for the mouse to fall from the rod were recorded for each trial. Mice were tested in 4 trials per day for 2 consecutive days or in 3 trials per day for 4 consecutive days. There was a 30–60-minute resting interval between trials.

#### Dowel test

A mouse was placed in the center of a horizontal dowel (6.5-mm or 9.5-mm diameter) and the latency to fall was measured with a maximal cutoff time of 120 seconds.

#### Inverted screen test

A mouse was placed onto a wire grid, and the grid was carefully picked up and shaken a couple of times to ensure that the mouse was holding on. The grid was then inverted such that the mouse was handing upside down from the grid. The latency to fall was measured with a maximal cutoff time of 60 seconds.

#### Wire hang test

A mouse was suspended by its forepaws on a 1.5-mm wire and the latency to fall was recorded with a maximal cutoff time of 60 seconds.

#### Foot slip test

A mouse was placed onto an elevated 40 × 25 cm wire grid (1 × 1 cm spacing) and allowed to freely move for 5 minutes. The number of foot slips was manually counted, and the moving distance was measured through a video camera (ANY-maze, Stoelting). The number of foot slips were normalized by the moving distance for each mouse.

#### Vertical pole test

A mouse was placed head-upward at the top of a vertical threaded metal pole (1.3-cm diameter, 55-cm length). The amount of time for the mouse to turn around and descend to the floor was measured with a maximal cutoff time of 120 seconds.

#### Grip strength

Forelimb grip strength was measured using a Grip Strength Meter (Columbus Instruments). A mouse was held by the tail and allowed to grasp a trapeze bar with its forepaws. Once the mouse grasped the bar with both paws, the mouse was pulled away from the bar until the bar was released. The digital meter displayed the level of tension exerted on the bar in gram-force (gf).

#### Acoustic startle response test

A mouse was placed in a well-ventilated, clear plastic cylinder and acclimated to the 70-dB background white noise for 5 minutes. The mouse was then tested with 4 blocks of 52 trials. Each block consisted of 13 trials, during which each of 13 different levels of sound (70, 74, 78, 82, 86, 90, 94, 98, 102, 106, 110, 114, or 118 dB, 40 milliseconds, inter-trial interval of 15 seconds on average) was presented in a pseudorandom order. The startle response was recorded for 40 milliseconds after the onset of the sound. The rapid force changes due to the startles were measured by an accelerometer (SR-LAB, San Diego Instruments).

#### Pre-pulse inhibition test

A mouse was placed in a well-ventilated, clear plastic cylinder and acclimated to the 70-dB background noise for 5 minutes. The mouse was then tested with 6 blocks of 48 trials. Each block consisted of 8 trials in a pseudorandom order: a “no stimulus” trial (40 milliseconds, only 70-dB background noise present), a test pulse trial (40 milliseconds, 120 dB), 3 different pre-pulse trials (20 milliseconds, 74, 78, or 82 dB), and 3 different pre-pulse inhibition trials (a 20-millisecond, 74, 78, or 82 dB pre-pulse preceding a 40-millisecond, 120-dB test pulse by 100 milliseconds). The startle response was recorded for 40 milliseconds after the onset of the 120-dB test pulse. The inter-trial interval is 15 seconds on average. The rapid force changes due to the startles were measured by an accelerometer (SR-LAB, San Diego Instruments). Pre-pulse inhibition of the startle responses was calculated as “1 – (pre-pulse inhibition trial/test pulse trial)”.

#### Hot plate test

A mouse was placed on a hot plate (Columbus Instruments) with a constant temperature of 55 °C. The latency for the mouse to first respond with either a hind paw lick, hind paw flick, or jump was measured. If the mouse did not respond within 45 seconds, then the test was terminated, and the latency was considered to be 45 seconds.

#### Novel object recognition test

A mouse was first habituated in an empty arena (24 × 45 × 20 cm) for 5 minutes before every trial. The habituated mouse was then placed into the testing arena with two identical objects for the first three trials. In the fourth trial, the mouse was exposed to the familiar object that it interacted during the previous three trials and a novel object. In the fifth trial, the mouse was presented with the two original, identical objects. Each trial lasted 5 minutes. The inter-trial interval was 24 hours or 5 minutes. In the modified version, *Stxbp1^tm1d/+^* and WT mice were exposed to the objects for 10 and 5 minutes during each trial, respectively. The movement of mice was recorded by a video camera placed above the test arena. The amount of time that the mouse interacted with the objects (*T*) was recorded using a wireless keyboard (ANY-maze, Stoelting). The preference index of interaction was calculated as *T*_novel object_/(*T*_familiarobject_ + *T*_novel object_).

#### Fear conditioning test

Pavlovian fear conditioning was conducted in a chamber (30 × 25 × 29 cm) that has a grid floor for delivering electrical shocks (Coulbourn Instruments). A camera above the chamber was used to monitor the mouse. During the 5-minute training phase, a mouse was placed in the chamber for 2 minutes, and then a sound (85 dB, white noise) was turned on for 30 seconds immediately followed by a mild foot shock (2 sec, 0.72 mA). The same sound and foot shock were repeated one more time 2 minutes after the first foot shock. After the second foot shock, the mouse stayed in the training chamber for at least 18 seconds before returning to its home cage. After 1 or 24 hours, the mouse was tested for the contextual and cued fear memories. In the contextual fear test, the mouse was placed in the same training chamber and its freezing behavior was monitored for 5 minutes without the sound stimulus. The mouse was then returned to its home cage. One to two hours later, the mouse was transferred to the chamber after it has been altered using plexiglass inserts and a different odor to create a new context for the cued fear test. After 3 minutes in the chamber, the same sound cue that was used in the training phase was turned on for 3 minutes without foot shocks while the freezing behavior was monitored. The freezing behavior was scored using an automated video-based system (FreezeFrame, Actimetrics). The freeze time (%) during the first 2 minutes of the training phase (i.e., before the first sound) was subtracted from the freeze time (%) during the contextual fear test. The freeze time (%) during the first 3 minutes of the cued fear test (i.e., without sound) was subtracted from the freeze time (%) during the last 3 minutes of the cued fear test (i.e., with sound).

#### Y maze spontaneous alternation test

A mouse was placed in the center of a Y-shaped maze consisting of three walled arms (35 × 5 × 10 cm) and allowed to freely explore the different arms for 10 minutes. The sequence of the arms that the mouse entered was recorded using a video camera (ANY-maze, Stoelting). The correct choice refers to when the mouse entered an alternate arm after it came out of one arm.

#### Elevated plus maze test

A mouse was placed in the center of an elevated maze consisting of two open arms (25 × 8 cm) and two closed arms with high black walls (25 × 8 × 15 cm) and allowed to freely explore for 10 minutes in the presence of 700–750-lux illumination and 65-dB background white noise. The mouse activity was recorded using a video camera (ANY-maze, Stoelting).

#### Light-dark chamber test

A mouse was placed in a rectangular light-dark chamber (44 × 21 × 21 cm) and allowed to freely explore for 10 minutes in the presence of 700–750 lux illumination and 65-dB background white noise. One third of the chamber is made of black plexiglass (dark) and two thirds is made of clear plexiglass (light) with a small opening between the two areas. The movement of the mouse was tracked by the Open Field Locomotor system (OmniTech).

#### Hole-board test

A mouse was placed at the center of a clear chamber (40 × 40 × 30 cm) that contains a black floor with 16 evenly spaced holes (5/8-inch diameter) arranged in a 4 × 4 array. The mouse was allowed to freely explore for 10 minutes. Its open-field activity above the floorboard and nose pokes into the holes Yes, were detected by infrared beams above and below the hole board using the VersaMax system (OmniTech).

#### Resident-intruder test

Male test mice (resident mice) were individually caged for 2 weeks before testing, and age-matched male white BALB/cAnNTac mice (Taconic) were group-housed to serve as the intruders. During the test, an intruder was placed into the home cage of a test mouse for 10 minutes, and their behaviors were video recorded. Videos were scored for the number and duration of each attack by the resident mouse regardless the attach was initiated by either the resident or intruder.

#### Tube test

A pair of a mutant mouse and an age- and sex-matched WT littermate that were housed in different home cages were placed into the opposite ends of a clear acrylic, cylindrical tube (3.5-cm diameter). The mouse that retreats backwards first was considered as the loser. The winner was scored as 1 and the loser as 0. Each mutant mouse was tested again 3 different WT littermates and the scores were averaged.

#### Three-chamber test

The apparatus (60.8 × 40.5 × 23 cm) consists of three chambers (left, center, and right) of equal size with 10 × 5 cm openings between the chambers. WT C57BL/6J mice were used as partner mice. A test mouse was placed in the apparatus with a mesh pencil cup in each of the left and right chambers and allowed to freely explore for 10 minutes. A novel object was then placed under one mesh pencil cup and an age- and sex-matched partner mouse under the other mesh pencil cup. The test mouse was allowed to freely explore for another 10 minutes. The position of the test mouse was tracked through a video camera (ANY-maze, Stoelting), and the approaches of the test mouse to the object or partner mouse were scored manually using a wireless keyboard. Partner mice were habituated to the mesh pencil cups in the apparatus for 1 hour per day for 2 days prior to testing. A partner mouse was used only in one test per day.

#### Partition test

The partitioned cage is a standard mouse cage (28.5 × 17.5 ×12 cm) divided in half with a clear perforated partition (a hole of 0.6-cm diameter). WT C57BL/6J mice were used as partner mice. A test mouse was housed in one side of the partitioned cage for overnight. In the afternoon before testing, an age- and sex-matched partner mouse was placed in the opposite half of the partitioned cage. On the next day, the time and number of approaches of the test mouse to the partition were scored using a handheld Psion event recorder (Observer, Noldus) in three 5-minute tests. The first test measured the approaches with the familiar overnight partner. The second measured the approaches with a novel partner mouse. The third test measured the approaches with the returned original partner mouse.

#### Nestlet shredding test

A mouse was individually housed in its home cage, and an autoclaved Nestlet was given to the mouse. The quality of the nest was assessed every 24 hours for 3 consecutive days.

#### Marble burying test

A clean standard housing cage was filled with approximately 8-cm deep bedding material. 20 marbles were arranged on top of the bedding in a 4 × 5 array. A mouse was placed into this cage and remained undisturbed for 30 minutes before returning to its home cage. The number of buried marbles (i.e., at least 2/3 of the marble covered by the bedding) was recorded.

### Video-EEG/EMG

Mice at 3–4 weeks of age were anesthetized with 2.5% isoflurane in oxygen, and the body temperature was maintained by a feedback based DC temperature control system at 37°C. The head was secured in a stereotaxic apparatus, and an incision was made along the midline to expose the skull. Craniotomies (approximate diameter of 0.25 mm) were performed with a round bur (0.25-mm diameter) and a high-speed rotary micromotor at coordinates (see below) that were normalized by the distance between Bregma and Lambda (DBL). Perfluoroalkoxy polymer (PFA)-coated silver wire electrodes (127-µm bare diameter, 177.8-µm coated diameter, A-M Systems) were used for grounding, referencing, and recording. A grounding electrode was placed on the right frontal cortex. An EEG reference electrode was placed on the cerebellum. Three EEG electrodes were placed on the left frontal cortex (anterior posterior (AP): 0.42 of DBL, medial lateral (ML): 0.356 of DBL, dorsal ventral (DV): −1.5 mm), left, and right somatosensory cortices (AP: −0.34 of DBL, ML: ± 0.653 of DBL, DV: −1.5mm). An EMG recording and an EMG reference electrode were inserted into the neck muscle. All the electrodes were connected to an adapter that was secured on the skull by dental acrylic. The skin around the wound was sutured, and mice were returned to the home cage to recover for at least one week. Before recording, mice were individually habituated in the recording chambers (10-inch diameter of plexiglass cylinder with bedding and access to food and water) for 24 hours. EEG/EMG signals were sampled at 5000 Hz with a 0.5-Hz high-pass filter, and synchronous videos were recorded at 30 frames per second from freely moving mice for continuous 72 hours using a 4-channel EEG/EMG tethered system (Pinnacle Technology).

To detect spike-wave discharges (SWDs), EEG signals of each channel were divided into 10-minute segments, and each segment was filtered by a third order Butterworth bandpass filter with 0.5–400 Hz cutoffs. The filtered data was divided into 250-millisecond non-overlapping epochs. EEG signal changes that occurred in the time domain were captured by root mean square (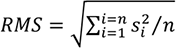; *s*, EEG signal; *n* = 1250), and spike density (number of spikes normalized to each epoch). EEG signal changes that occurred in the frequency domain were captured by frequency band ratio 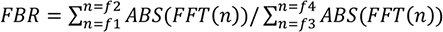; *f1* = 100; *f2* = 300; *f3* = 0.5; *f4* = 80) where the power of the upper band (100–300 Hz) was contrasted with that of the lower band (0.5–80 Hz). The above features were computed in MATLAB. An EEG segment that exceeded thresholds for all of the above features was identified as a SWD candidate. The candidates were further classified by a convolutional neural network in Python that was trained with manually labeled EEG segments. The first layer of the network contained 32 filters that returned their matches with 10-millisecond (kernel size) non-overlapping (stride) candidate segments across the three EEG channels. Successive convolutional layers were stacked sequentially. For every two consecutive convolutional layers, there was a pooling layer that down-sampled the outputs by a factor of 5 to reduce computation. The overall network consisted of two layers of 32 filters, one layer of pooling, two layers of 64 filters, one layer of pooling, two layers of 128 filters, and one layer of pooling. The network was trained through an iterative approach. In each training iteration, the optimizer (Adadelta) updated the weights of the filters, and the loss function (binary cross entropy) evaluated how well the network predicted SWDs. This iteration process continued until the loss function was minimized. Methods implemented to reduce overfitting included dropout (i.e., 50% of the neurons were randomly dropped out from calculation for each iteration) and early stopping (i.e., training process was stopped when the loss function on validation set did not decrease for 3 iterations). The trained neural network removed 99% of the false-positive candidates and the remaining candidates were further confirmed by visual inspection. For each SWD, the duration (the time difference between the first and last peaks) and spike rate were quantified. The SWD cluster was defined as a cluster of 5 or more SWD episodes that occurred with inter-episode-interval of maximal 60 s.

To identify myoclonic seizures, we visually inspected the EEG/EMG traces and videos to identify sudden jumps and myoclonic jerks. When the mouse suddenly and quickly move the body in less than one second, if one or more limbs leave the cage floor, then this is classified as a sudden jump. If all limbs stay on the cage floor, then this is classified as a myoclonic jerk. The state of the mouse right before the myoclonic seizure was classified as REM sleep, NREM sleep, or awake based on the EEG/EMG.

### Brain slice electrophysiology

Mice were anesthetized by an intraperitoneal injection of a ketamine and xylazine mix (80 mg/kg and 16 mg/kg, respectively) and transcardially perfused with cold (0–4°C) slice cutting solution containing 80 mM NaCl, 2.5 mM KCl, 1.3 mM NaH_2_PO_4_, 26 mM NaHCO_3_, 4 mM MgCl_2_, 0.5 mM CaCl_2_, 20 mM D-glucose, 75 mM sucrose and 0.5 mM sodium ascorbate (315 mosmol, pH 7.4, saturated with 95% O_2_/5% CO_2_). Brains were removed and sectioned in the cutting solution with a VT1200S vibratome (Leica) to obtain 300-µm coronal slices. Slices were incubated in a custom-made interface holding chamber saturated with 95% O_2_/5% CO_2_ at 34 °C for 30 min and then at room temperature for 20 minutes to 8 hours until they were transferred to the recording chamber.

Recordings were performed on submerged slices in artificial cerebrospinal fluid (ACSF) containing 119 mM NaCl, 2.5 mM KCl, 1.3 mM NaH_2_PO_4_, 26 mM NaHCO_3_, 1.3 mM MgCl_2_, 2.5 mM CaCl_2_, 20 mM D-glucose and 0.5 mM sodium ascorbate (305 mosmol, pH 7.4, saturated with 95% O_2_/5% CO_2_, perfused at 3 ml/min) at 30–32°C. For whole-cell recordings, we used a K^+^-based pipette solution containing 142 mM K^+^-gluconate, 10 mM HEPES, 1 mM EGTA, 2.5 mM MgCl_2_, 4 mM ATP-Mg, 0.3 mM GTP-Na, 10 mM Na_2_-phosphocreatine (295 mosmol, pH 7.35) or a Cs^+^-based pipette solution containing 121 mM Cs^+^-methanesulfonate, 1.5 mM MgCl_2_, 10 mM HEPES, 10 mM EGTA, 4 mM Mg-ATP, 0.3 mM Na-GTP, 10 mM Na_2_-Phosphocreatine, and 2 mM QX314-Cl (295 mosmol, pH 7.35). Membrane potentials were not corrected for liquid junction potential (experimentally measured as 12.5 mV for the K^+^-based pipette solution and 9.5 mV for the Cs^+^-based pipette solution).

Neurons were visualized with video-assisted infrared differential interference contrast imaging and fluorescent neurons were identified by epifluorescence imaging under a water immersion objective (40×, 0.8 numerical aperture) on an upright SliceScope Pro 1000 microscope (Scientifica) with an infrared IR-1000 CCD camera (DAGE-MTI). Data were low-pass filtered at 4 kHz and acquired at 10 kHz with an Axon Multiclamp 700B amplifier and an Axon Digidata 1550 or 1440 Data Acquisition System under the control of Clampex 10.7 (Molecular Devices). Data were analyzed offline using AxoGraph X (AxoGraph Scientific).

Neuronal intrinsic excitability was examined with the K^+^-based pipette solution. The resting membrane potential was recorded in the whole-cell current clamp mode within the first minute after break-in. After balancing the bridge, the input resistance was measured by injecting a 500-ms hyperpolarizing current pulse (10–100 pA) to generate a small membrane potential hyperpolarization (2–10 mV) from the resting membrane potential. Depolarizing currents were increased in 5- or 10-pA steps to identify rheobase currents.

Postsynaptic currents were recorded in the whole-cell voltage clamp mode with the Cs^+^-based patch pipette solution. Only recordings with series resistance below 20 MΩ were included. IPSCs were recorded at the reversal potential for excitation (+10 mV). To record unitary connections between inhibitory interneurons and pyramidal neurons, Pv and Sst interneurons were identified by the Cre-dependent expression of tdTomato. Pyramidal neurons were first recorded in whole-cell voltage clamp mode (+10 mV) with the Cs^+^-based patch pipette solution, and a nearby Pv or Sst interneuron was subsequently recorded in the whole-cell current clamp mode with the K^+^-based patch pipette solution. Action potentials were elicited in Pv or Sst interneurons by a 2-ms depolarizing current step (1–2 nA) with 10 s inter-stimulus intervals. Unitary IPSC (uIPSC) amplitudes were measured from the average of 30–50 sweeps. We considered a Pvalb or Sst interneuron to be connected with a pyramidal neuron when the average uIPSC amplitude was at least three times the baseline standard deviation.

### Statistics

All reported sample numbers (*n*) represent biological replicates that are the numbers of tested mice or recorded neurons. Statistical analyses were performed with Prism 6 (GraphPad Software). D’Agostino-Pearson, Shapiro-Wilk, and Kolmogorov-Smirnov tests were used to determine if data were normally distributed. If all data within one experiment passed all three normality tests, then the statistical test that assumes a Gaussian distribution was used. Otherwise, the statistical test that assumes a non-Gaussian distribution was used. All statistical tests were two-tailed with an alpha of 0.05. The details of all statistical tests, numbers of replicates and mice, and *P* values were reported in ***Supplementary Table 2*.**

## AUTHOR CONTRIBUTIONS

M.X. and W.C. designed the study, reviewed and interpreted the data. H.C., H.T.C., and M.X. generated the transgenic mice and performed the initial characterizations of transgenic mice. W.C. performed and analyzed the biochemical, behavioral and EEG experiments. Z.L.C. and J.E.M. performed the slice electrophysiology experiments. E.S.C and J.H.K. contributed to the EEG data analysis. S.H. and J.T. contributed to the initial EEG experiments. H.Y.Z. and J.W.S. supervised the generation of transgenic mice. M.X. supervised all experiments. M.X. and W.C. wrote the manuscript with inputs from all authors.

## ACKNOWLEDGMENTS

This article is dedicated to the memory of Caroline DeLuca, who had *STXBP1* encephalopathy and inspired this project. We thank Gabriele Schuster for ES cell work and blastocyst injection, Corinne Spencer and James Frost for suggestions and discussions. This work was supported in part by Citizens United for Research in Epilepsy (to M.X.), the National Institute of Neurological Disorders and Stroke (R01NS100893 to M.X.), American Epilepsy Society Postdoctoral Research Fellowship (to W.C.), the Eunice Kennedy Shriver National Institute of Child Health and Human Development (U54HD083092 to Baylor College of Medicine Intellectual and Developmental Disabilities Research Center, Neurobehavioral Core), and the In Vivo Neurophysiology Core of Jan and Dan Duncan Neurological Research Institute. H.Y.Z. is a Howard Hughes Medical Institute investigator. M.X. is a Caroline DeLuca Scholar.

---

**Figure 1-supplement 1.**
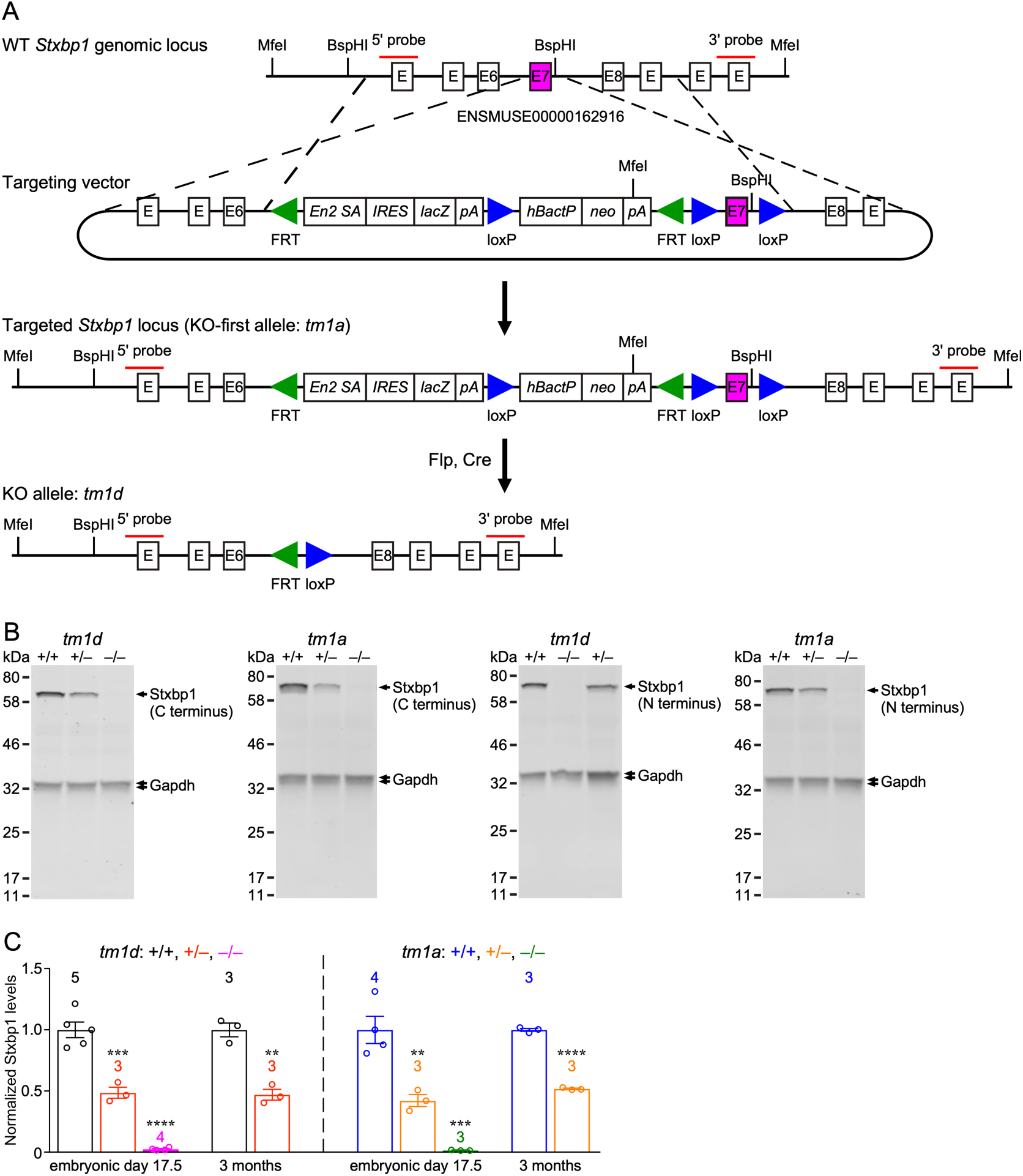
Generation of two new *Stxbp1* null alleles. (**A**) The *Stxbp1* WT genomic region was targeted by a multipurpose cassette that contains an *Engrailed 2* splice acceptor site (*En2SA*), an encephalomyocarditis virus internal ribosomal entry site (*IRES*), *lacZ*, SV40 polyadenylation element (pA), and floxed exon 7, resulting in the KO-first allele (*tm1a*). The restriction enzymes and probes used in the Southern blots are indicated in the diagrams. The KO-first allele was converted to the KO allele (*tm1d*) by crossing *Stxbp1^tm1a/+^* mice with *Rosa26-Flpo* and *Sox2-Cre* mice sequentially. (**B**) Representative Western blots of Stxbp1 and Gapdh proteins extracted from the brains at embryonic day 17.5. Stxbp1 was detected by an antibody recognizing the C terminus (left two blots) or the N terminus (right two blots). The genotypes are indicated above the samples. Note that Stxbp1 was reduced in heterozygous mutants and absent in homozygous mutants. (**C**) Summary data of normalized Stxbp1 expression levels at the ages of embryonic day 17.5 and 3 months. Stxbp1 levels were first normalized by the Gapdh levels to obtain the relative expression levels of Stxbp1. The relative expression levels of Stxbp1 were then normalized by the average Stxbp1 levels of all WT mice from the same blot. The data obtained by both Stxbp1 antibodies are combined. The numbers of analyzed mice are indicated in the figures. Each circle represents one mouse. Bar graphs are mean ± s.e.m. ** *P* < 0.01, *** *P* < 0.001, **** *P* < 0.0001.

**Figure 1-supplement 2.**
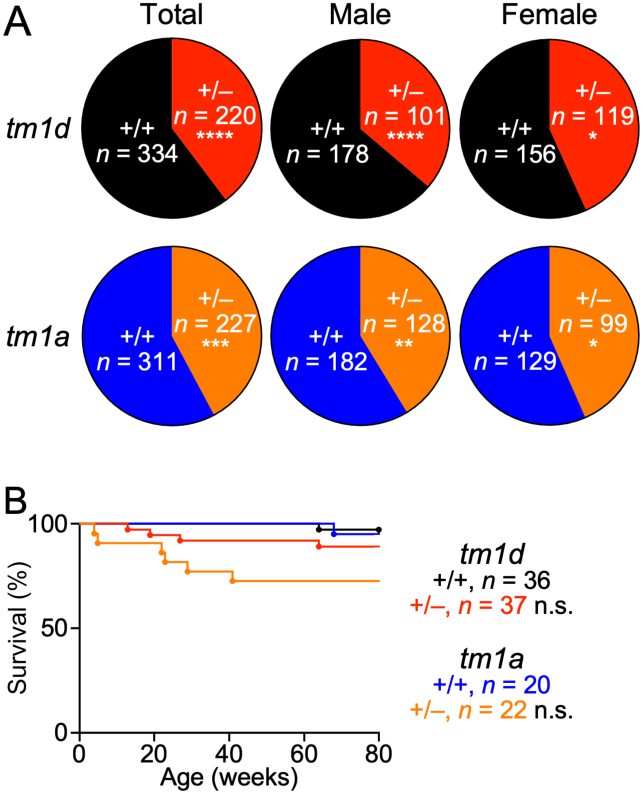
Reduced survival of *Stxbp1* haploinsufficient mice. (**A**) *Stxbp1^tm1d/+^* and *Stxbp1^tm1a/+^* male mice were crossed with WT female mice. The observed genotypes of the offspring at weaning (i.e., around the age of 3 weeks) are shown in the pie charts. The male, female, and total *Stxbp1^tm1d/+^* and *Stxbp1^tm1a/+^* mice were significantly less than Mendelian expectations. Note that the genotypes of some female mice were not determined and therefore, they were not included in this analysis. (**B**) Survival curves of a subset of *Stxbp1^tm1d/+^*, *Stxbp1^tm1a/+^*, and WT mice that were monitored for 80 weeks. The numbers of observed mice are indicated in the figures. n.s. *P* > 0.05, ** *P* < 0.01, *** *P* < 0.001, **** *P* < 0.0001.

**Figure 1-supplement 3.**
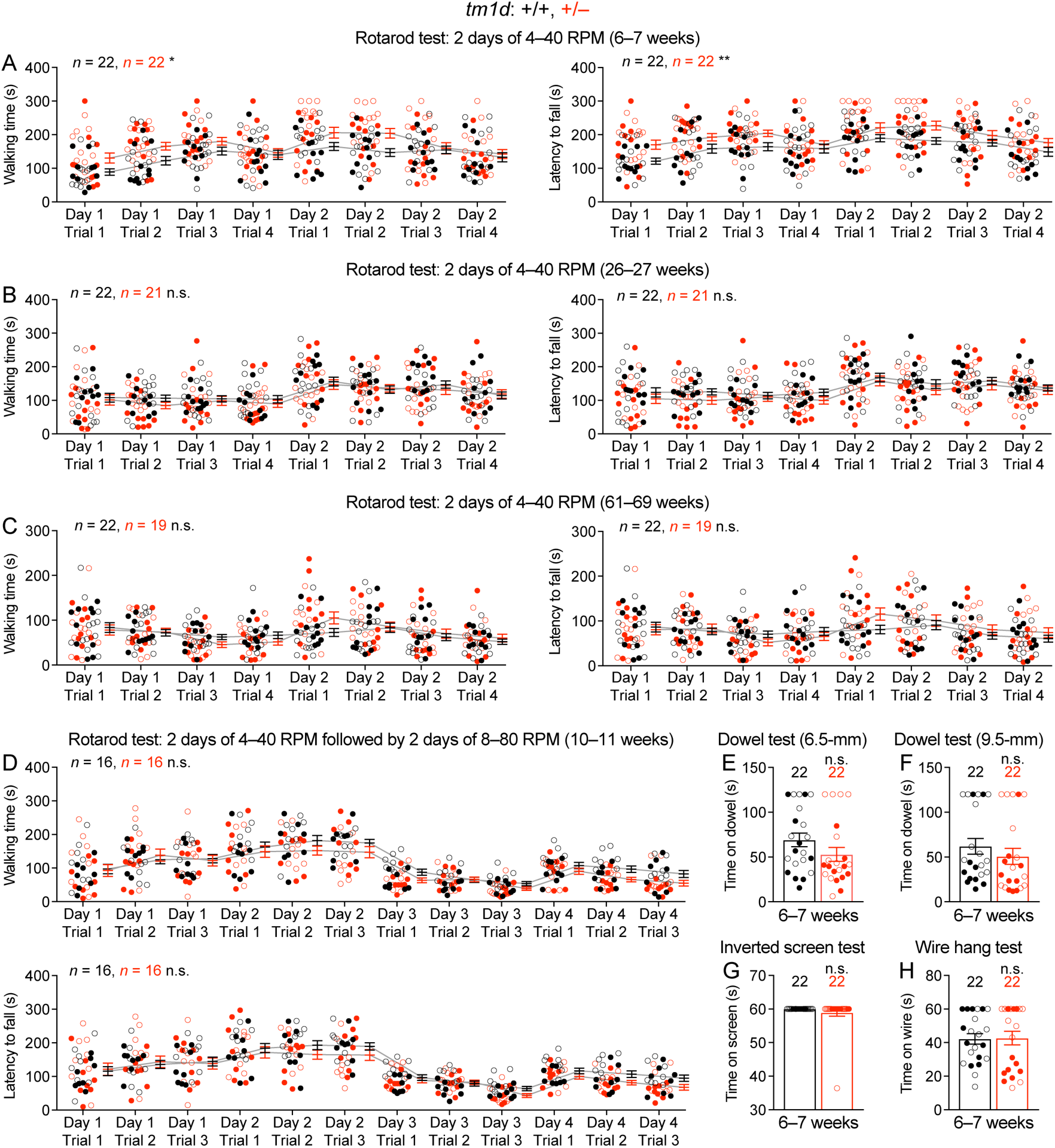
Normal performance of *Stxbp1^tm1d/+^* mice in rotarod, dowel, inverted screen, and wire hang tests. (**A**) In the 2-day rotarod test, 6–7-week old *Stxbp1^tm1d/+^* mice performed better than WT mice, as they were able to walk (left panel) and stay (right panel) on the rotating rod for longer time, probably due to their lower body weights or hyperactivity. (**B,C**) Similar to (A), but for the ages of 26–27 weeks (B) and 61–69 weeks (C). *Stxbp1^tm1d/+^* mice performed similar to WT mice. (**D**) In the 4-day rotarod test, *Stxbp1^tm1d/+^* mice performed similar to WT mice at the age of 10–11 weeks. (**E,F**) *Stxbp1^tm1d/+^* mice could stay on the dowel (6.5- or 9.5-mm diameter) for similar amount of time as WT mice. (**G,H**) *Stxbp1^tm1d/+^* mice could hang on the screen (G) or wire (H) for similar amount of time as WT mice. The numbers and ages of tested mice are indicated in the figures. Each filled (male) or open (female) circle represents one mouse. Bar graphs are mean ± s.e.m. n.s. *P* > 0.05, * *P* < 0.05, ** *P* < 0.01.

**Figure 1-supplement 4.**
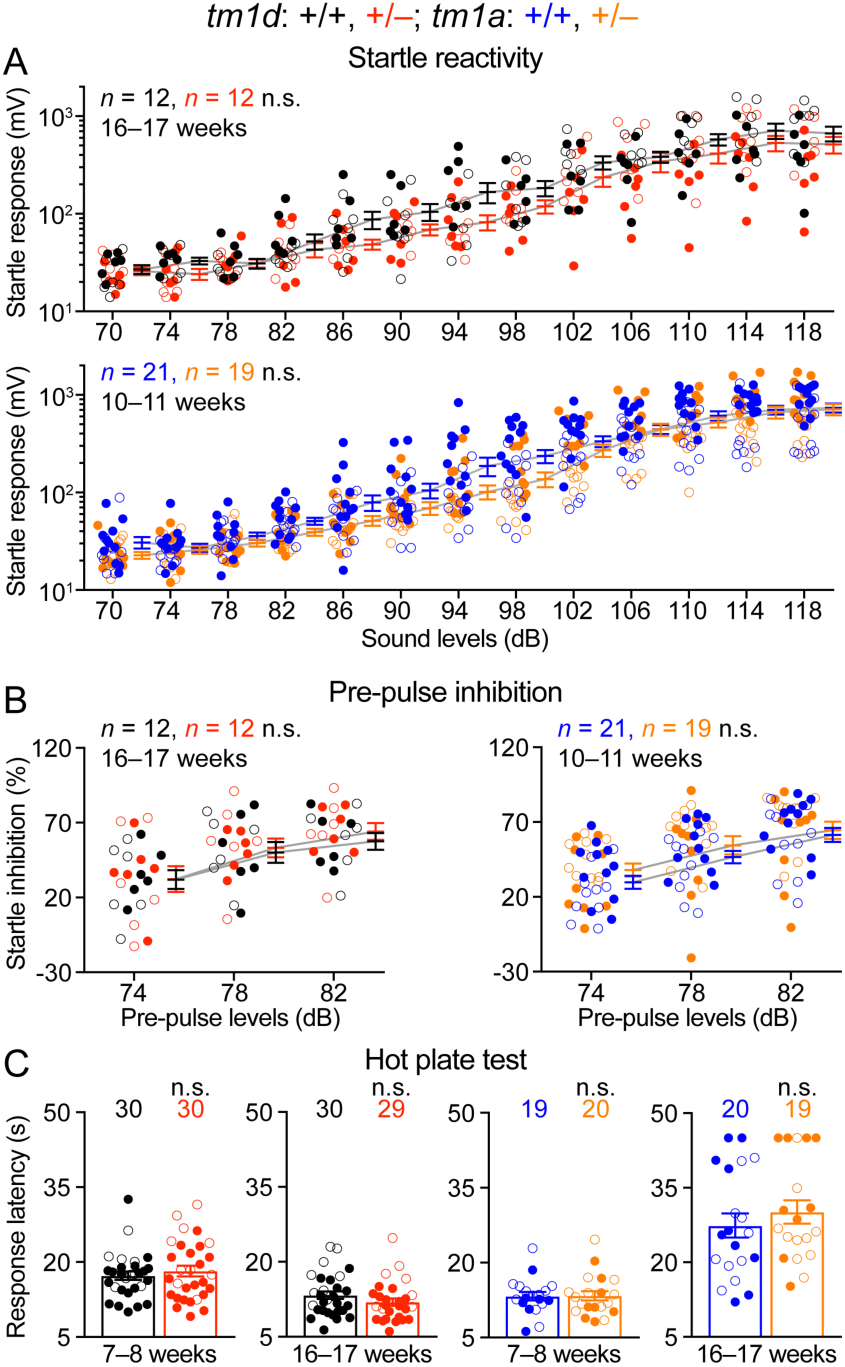
*Stxbp1* haploinsufficient mice have normal sensory functions. (**A**) *Stxbp1^tm1d/+^* and *Stxbp1^tm1a/+^* mice showed similar acoustic startle responses as WT mice at different sound levels. (**B**) In the pre-pulse inhibition test, when a weak sound (74, 78, or 82 dB) preceded a loud sound (120 dB), *Stxbp1^tm1d/+^* and *Stxbp1^tm1a/+^* mice showed a similar reduction in the startle responses to the loud sound as WT mice. (**C**) In the hot plate test, *Stxbp1^tm1d/+^*and *Stxbp1^tm1a/+^* mice showed similar latencies in response to the high temperature as WT mice. The numbers and ages of tested mice are indicated in the figures. Each filled (male) or open (female) circle represents one mouse. Bar graphs are mean ± s.e.m. n.s. *P* > 0.05.

**Figure 2-supplement 1.**
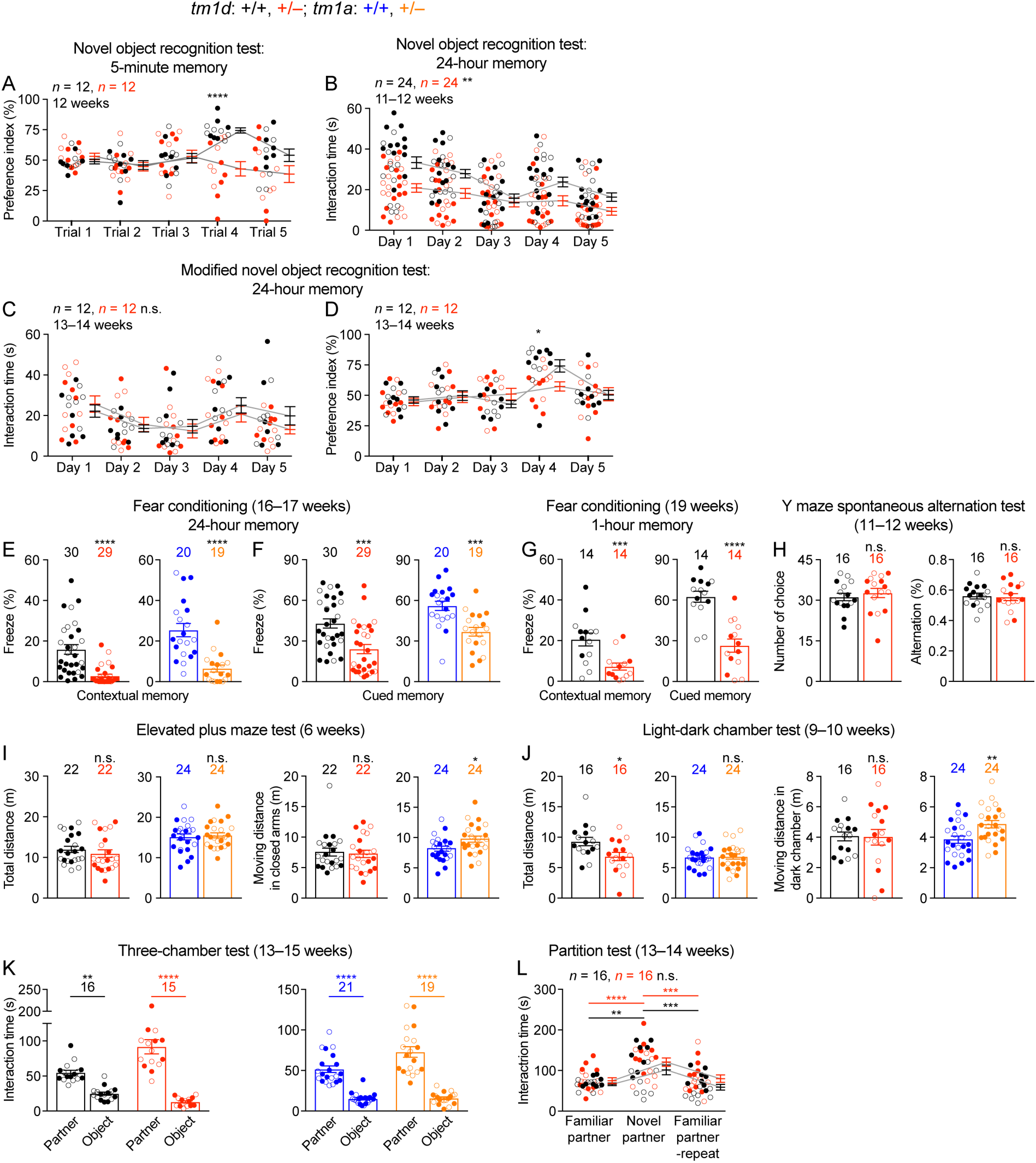
*Stxbp1* haploinsufficient mice show impaired cognition and normal social interactions. (**A**) In the novel object recognition test with 5-minute testing intervals, *Stxbp1^tm1d/+^* mice did not show a preference for the novel object on trial 4 when they were presented with a familiar and a novel object. (**B**) In the novel object recognition test with 24-hour testing intervals (same as Figure 2A), *Stxbp1^tm1d/+^* mice spent less time interacting with the familiar and novel objects. (**C,D**) In the modified novel object recognition test with 24-hour testing intervals, *Stxbp1^tm1d/+^* mice spent similar amount of time interacting with the familiar and novel objects as WT mice (C), but they still failed to show a preference for the novel object on day 4 (D). (**E,F**) At the age of 16–17 weeks, *Stxbp1^tm1d/+^*and *Stxbp1^tm1a/+^* mice showed a reduction in both contextual (E) and cued (F) fear memories 24 hours after training. (**G**) *Stxbp1^tm1d/+^* mice showed a reduction in both contextual (left panel) and cued (right panel) fear memories 1 hour after training. (**H**) In the Y maze spontaneous alternation test that evaluates working memory, *Stxbp1^tm1d/+^* mice made similar numbers of choices (left panel) and alternations (right panel) as WT mice. (**I**) In the elevated plus maze test, the total travel distances and travel distances in the closed arms of *Stxbp1^tm1d/+^*and *Stxbp1^tm1a/+^* mice were similar or slightly longer than those of WT mice. (**J**) In the light-dark chamber test, the total travel distances of *Stxbp1^tm1d/+^* mice were reduced due to the reduction of their travel distances in the light chamber and normal travel distances in the dark chamber. The total travel distances *Stxbp1^tm1a/+^* mice were normal and their travel distances in the dark chamber was slightly increased as compared to WT mice. (**K**) In the three-chamber test, *Stxbp1^tm1d/+^* and *Stxbp1^tm1a/+^* mice showed a preference in interacting with the partner mouse over the object. (**L**) In the partition test, *Stxbp1^tm1d/+^* mice showed a normal preference for the novel partner mouse. The numbers and ages of tested mice are indicated in the figures. Each filled (male) or open (female) circle represents one mouse. Bar graphs are mean ± s.e.m. n.s. *P* > 0.05, * *P* < 0.05, ** *P* < 0.01, *** *P* < 0.001, **** *P* < 0.0001.

**Figure 3-supplement 1.**
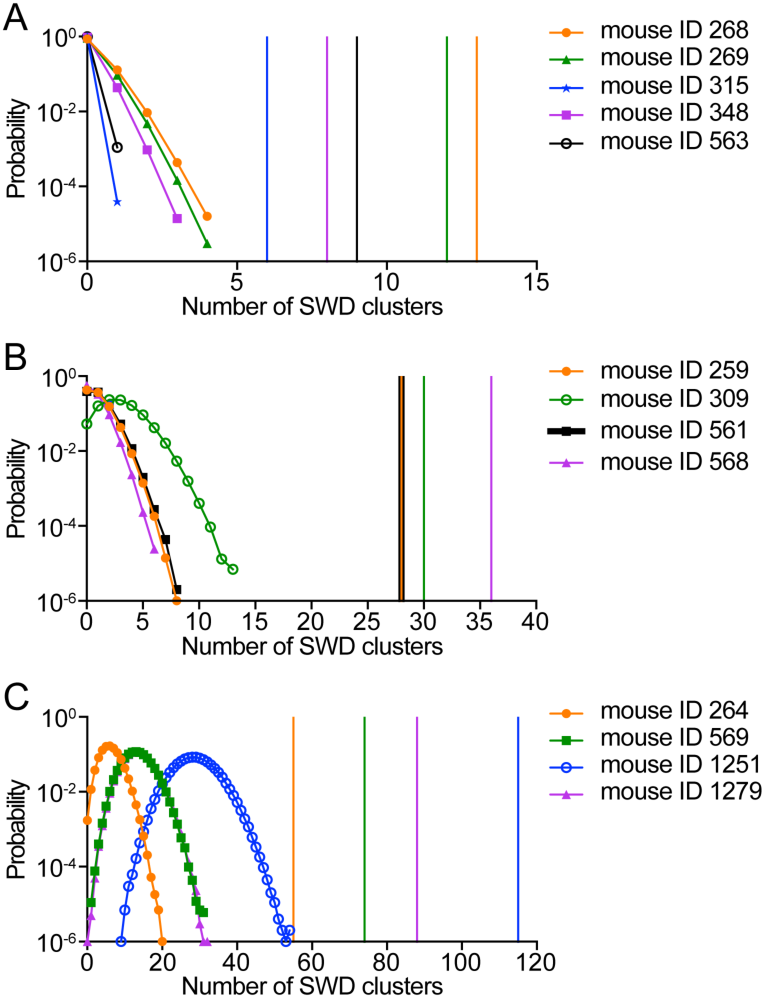
The clustering of SWDs in *Stxbp1^tm1d/+^* mice does not result from a random distribution of frequent SWD episodes. (**A–C**) In *Stxbp1^tm1d/+^* mice, many SWDs occurred in a cluster manner. A SWD cluster is defined as 5 or more episodes of SWDs that occur with an inter-episode-interval of 60 s or less. For each *Stxbp1^tm1d/+^* mouse, simulations were performed to determine if the clustering of SWD episodes was simply due to the overall high frequencies of episodes. The recorded episodes of SWDs from a *Stxbp1^tm1d/+^*mouse were randomly distributed in the same period of time for 10^6^ times. The number of SWD clusters was determined from each simulated distribution, and the results of the 10^6^ simulations are shown as the probability distribution of the number of SWD clusters for each mouse. The vertical lines with the same color as the probability distribution curves represent the numbers of the recorded SWD clusters in each mouse. The numbers of simulated SWD clusters are all smaller than that of recorded SWD clusters for each *Stxbp1^tm1d/+^* mouse (*P* < 10^-6^), demonstrating that a random distribution of the same number of SWD episodes does not result in the same clustering of SWDs in *Stxbp1^tm1d/+^* mice.

**Figure 3-supplement 2–5. Video-EEG/EMG recordings from *Stxbp1^tm1d/+^* mice.**

Representative videos showing a SWD cluster (***Video S1***), a long SWD during REM sleep (***Video S2***), a myoclonic jump (***Video S3***), and a myoclonic jerk (***Video S4***). The EEG/EMG traces (from top to bottom) were from the left frontal cortex, right somatosensory cortex, left somatosensory cortex, and neck muscle. The vertical line indicates the time of the current video frame. Note that the EEG signal from the left somatosensory cortex (the third channel) is inverted.

**Figure 4-supplement 1.**
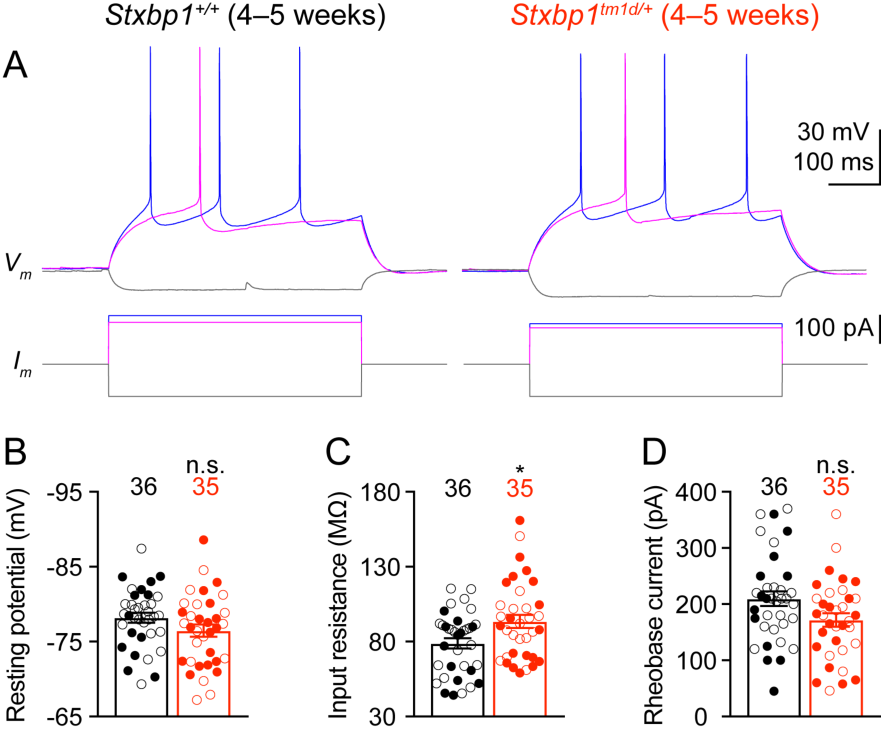
Intrinsic neuronal excitability of *Stxbp1^tm1d/+^* mice is slightly increased. (**A**) Membrane potentials (upper panels) in response to current injections (lower panels) in layer 2/3 pyramidal neurons of the somatosensory cortex from WT and *Stxbp1^tm1d/+^* mice. (**B–D**) Summary data showing that *Stxbp1^tm1d/+^*neurons had similar resting membrane potentials and rheobase currents as WT neurons, but their input resistances were 19% larger than WT neurons. The numbers of recorded neurons are indicated in the figures. Each filled (male) or open (female) circle represents one neuron. Bar graphs are mean ± s.e.m. n.s. *P* > 0.05, * *P* < 0.05.

